# Deficient Memory, Long-Term Potentiation and Hippocampal Synaptic Plasticity in Galectin-4-Deficient Mice

**DOI:** 10.1101/2024.10.14.617868

**Authors:** María Elvira Brocca, Arancha Mora-Rubio, Elena Alonso-Calviño, Elena Fernández-López, Natalia Díez-Revuelta, Juan C. Guirado, Rebeca Álvarez-Vélez, Ana Dopazo, Esperanza López-Merino, David Martos-Puñal, José A. Esteban, Juan Aguilar, Alonso M. Higuero, José Abad-Rodríguez

## Abstract

**BACKGROUND:** Brain function is influenced by the gut through the microbiota-gut-brain axis. Non-physiological microbiota-depletion or induced gut infection in animal models, have been instrumental to link intestinal alterations to cognitive and mood dysfunctions. However, the effects of specific, controlled, physiologically relevant shifts in commensal microbiota composition on brain function remain poorly understood.

**METHODS:** Mice deficient in galectin-4 (Lgals4-KO) were used in this study. Gut microbiota was analysed by 16S-rRNA sequencing. Cognitive and mood status were evaluated with specific behavioral tests. Long-term potentiation (LTP) was tested *ex vivo* and *in vivo* by electrophysiological methods and *in vitro* by immunofluorescence and western blot. RNA-sequencing was used for transcriptomic analyses. Golgi-Cox staining and transmission electron microscopy were used for quantitative and morphological assessments of dendritic spines and synapses.

**RESULTS:** Lgals4-KO mice present an altered intestinal commensal microbiota in the absence of pathogens, deficient memory formation, and impaired hippocampal LTP *in vivo* and *ex vivo*. Furthermore, Lgals4-KO neurons show a reduced activation of AMPA receptors and of CaMKII upon chemically induced LTP *in vitro*. These mice also display significantly lower dendritic spine density and shorter spine length in hippocampal dendrites, as well as an increased area of the postsynaptic densities

**CONCLUSIONS:** Our results define a new role for galectin-4 in the modulation of commensal bacteria. We also show that the absence of galectin-4 induces changes in gut microbial composition, along with synaptic alterations and memory impairment, supporting our hypothesis that variations in endogenous microbiota may cause or contribute to relevant neurological pathologies.

## INTRODUCTION

Brain function and many of its pathological conditions are influenced by biochemical or nerve signals that, originated in the gut, reach the central nervous system through the microbiota-gut-brain axis (1–3). Within the gut, epithelial glycans are a major component of the intestinal mucosa (4–6) that regulate the interactions between the intestinal epithelium and the commensal or pathogenic bacteria, whose surfaces are also heavily glycosylated (7). Alterations of glycan-based interactions can disrupt mucosal barrier function facilitating pathogenic intestinal infections (8–11). Associated gut inflammatory or infectious processes, as well as the extreme depletion of microbiota in germ-free or antibiotic-treated animal models, have been linked to learning, memory and mood dysfunctions, and other neurodegenerative pathologies (2,12,13). However, the impact of controlled, physiologically relevant shifts in commensal microbiota composition on specific brain functions remains largely undefined.

In the intestinal lumen, pathogenic bacteria that express glycosylated host-like antigens can evade detection by the adaptive immune system through molecular mimicry. This evasion enables them to invade the intestinal epithelium, triggering intestinal dysfunctions and immunological reactions (22). Galectin-4 (Gal-4), a lectin prominently expressed by epithelial cells in the intestinal tract (20,21), cross-links glycosylated molecules containing β-galactoside moieties (16), O-linked sulfoglycans or sulphated lipids (17–19). Through specific interactions with glycosylated self-antigens, Gal-4 targets and eliminates these pathogenic bacteria, either extracellularly, as reported for *Escherichia coli* (23–25), or intracellularly in epithelial cells infected with *Salmonella enterica* (26). Despite its activity in the control of intestinal pathogenic bacteria, no role of Gal-4 in commensal bacteria regulation had been reported so far.

We leveraged the use of a Gal-4-deficient mouse model (Lgals4-KO) as a novel strategy to evaluate the effect of endogenous microbiota compositional alterations on cognitive functions.

## METHODS AND MATERIALS

See Supplemental Methods for detailed methodology.

### Animals

The Gal-4-deficient mice (C57BL/6NJ-Lgals4em1(IMPC)J/J), were obtained from The Jackson Laboratories. C57BL/6NJ mice were used as controls. All experimental procedures involving animals were approved by the Animal Bioethics Committee (CEBA) of the Hospital Nacional de Parapléjicos-Research Center.

### Intestinal microbiota analysis

DNA was extracted from stool samples using QIAamp Fast DNA Stool Mini Kit (Qiagen). The V3-V4 hypervariable region of the 16S rRNA gene was amplified barcoded and sequenced on an Illumina MiSeq platform. The raw sequencing reads were processed using DADA2 (v.1.32.0) in RStudio (v.2024.04.2+764, R v.4.4.0), and taxonomy was assigned using SILVA (version 138.1) and NCBI reference datasets. Alpha and beta diversities were assessed using standard indexes. Differential microbial abundance was evaluated using ANCOM-BC, (v.2.6.0) and LEfSe.

### Behavior tests

Sessions were video recorded with automatic tracking. *Y-Maze:* Mice were trained for 5 min with one arm closed, followed 30 min later by a 5 min test session allowing free exploration of all arms. Arm entries, latencies and time spent in each arm were recorded. *Novel Object Recognition (NOR) and Object Location (OL) tasks:* Mice freely explored the square open arena for 15 min. The day after animals performed 2×10 min training sessions exploring two similar objects made from different materials in adjacent corners of the arena. For NOR test, one object was replaced with a novel object in the same location. For the OL test, one object was moved to the opposite corner. In both cases, data from 10 min exploration periods were analysed. The *Tail Suspension* and *the Forced Swim tests* were performed as previously described (27,28). For the *Elevated Zero Maze test* mice were placed at the boundary between an open and a closed zone of the maze, facing the closed zone, and recorded during a 5 min exploration period. The time spent in the open zones was analysed.

### RT-PCR and RNA-seq

Total RNA from hippocampi (30-day-old mice) was extracted (RNeasy Mini Kit, Qiagen). *RT-PCR:* cDNA was synthesized with High-Capacity RNA-to-cDNA kit (Applied Biosystems). Expression was measured by RT-PCR (7900HT system, Applied Biosystems) using TaqMan probes (Applied Biosystems, Supplemental Table S2). GAPDH and β-actin served as controls. The relative amounts of mRNA were calculated using the 2^(−ΔΔCT)^ method (29). *RNA-seq: Libraries were prepared from* 100 ng of total RNA (NEBNext Ultra II kit, New England Biolabs) and sequenced on a HiSeq 4000 (Illumina). Differential expression analysis was conducted using DESeq2 (version 1.44.0) (30), with adjusted p-values (P_adj_)< 0.05 and log_2_FC ≥ 0.1 thresholds. Functional classification relied on Gene Ontology (GO) annotations (GO.db version 3.19.1).

### Neuron cultures and cLTP induction

Primary mouse hippocampal neuron cultures were obtained from P17 embryos as previously reported (31) and maintained at 37°C and 5% CO_2_. Cells, previously exposed or not to recombinant human Gal-4 (0.75 mg/ml, 2 hours), were subjected to 15 min chemical LTP induction (Mg^2+^-free artificial cerebrospinal fluid -aCSF-, 0.1 μM rolipram, 50 μM forskolin and 100 μM picrotoxin). Neurons were then processed for immunocytochemistry or biochemical analyses.

### Electrophysiology

#### Brain slices (ex vivo)

Coronal brain slices (350 μm) were perfused with carbogenated aCSF, 100 μM picrotoxin. Field excitatory postsynaptic potentials (fEPSPs) were recorded in CA1 by stimulating Schaffer collateral fibers (50 μs pulses, 20-250 μA). Data were acquired with MultiClamp 700A/B amplifiers and pClamp software (Molecular Devices).

#### Living animal (in vivo)

Isofluorane-anesthetized animals underwent electrode placement in CA1 (pyramidal layer) and contralateral CA3. Local field potentials (LFPs, unfiltered) were recorded in CA1 in response to CA3 stimulation, and response magnitudes of the populational spikes were quantified across the entire stimulation protocol. Data were normalized to the average population response. Paired t-tests assessed intra-group differences (baseline vs. post-LTP) and inter-group comparisons.

### Histochemistry and Immunohistochemistry

#### Tissue sampling

Mice were anesthetized, perfused, and brains and intestines were collected. For immunohistochemistry (IHC) and Black Gold staining, tissues were post-fixed and then either cryoprotected (brains) or paraffin-embedded (intestines) before sectioning (brains: 40 μm; intestines: 4 μm). For pCamkII immunofluorescence, brains from mice subjected or not to LTP induction were post-fixed, cryoprotected, frozen and cryosectioned (20 μm). Dendritic spines were studied using Golgi-Cox staining on 100 μm brain sections from 7-month-old mice.

#### Immunostainings

For immunofluorescence, sections were deparaffinized, blocked and sequentially incubated with the indicated primary and fluorophore-conjugated secondary antibodies, and counterstained with DAPI. For immunohistochemical staining endogenous peroxidase activity was blocked, followed by incubation with primary antibodies and biotinylated secondary antibodies (Supplemental Table S1). Staining was visualized with Vectastain Elite ABC Kit Peroxidase (Vector). Black Gold *and* Golgi-Cox stainings were performed using the Black Gold II Myelin staining kit (#AG105, Millipore), and the FD Rapid GolgiStain™ Kit (FD NeuroTechnologies), respectively, according to manufacturers’ instructions. Detailed protocols for staining and image acquisition, segmentation and analysis are available in Supplemental Information and in (32).

### Transmission electron microscopy

Brains from 8-month-old mice were perfused, sectioned (200 μm, dorsal CA1), postfixed with 1% osmium tetroxide, 0.8% aqueous potassium ferricyanide, and sequentially treated with 0.15% tannic acid and 2% uranyl acetate. Upon dehydration, samples were embedded in epoxy resin. Serial ultrathin sections (70 nm) were stained with 2% uranyl acetate and lead citrate. Imaging was performed using a JEM-1400 Flash transmission electron microscope (Jeol, Japan) at 100 kV, with a Oneview 4K x4K CMOS camera (Gatan, Pleasanton, CA USA). Synaptic density and post-synaptic density dimensions in the *stratum radiatum* of CA1 were quantified using the unbiased physical dissector method (33), in combination with deep learning and manual annotation.

## RESULTS

### Lgals4-KO mice present an altered intestinal microbiota composition compared to WT animals

Gut microbiota composition was analysed and compared between Lgals4-KO and WT mice. Bacterial microbiota diversity (beta diversity) showed significant differences between mouse strains across all applied metrics (Bray-Curtis distance, Jaccard index, and unweighted UniFrac index) as determined by PERMANOVA (Adonis test; p=0.001) (Fig.1A, Fig.S1A-B). In contrast, the diversity within individual samples (alpha diversity) showed no differences (Fig.S1C). ANCOM-BC analysis identified significant reductions in Lgals4-KO mice for the genera *Peptococcus, UCG-009* and an unspecified genus of the *Prevotellaceae* family, while the genus *Paludicola* and the species *Alistipes_timonensis* were significantly increased (Figs.1B and S2A). Consistently, LEfSe analysis revealed seven significantly increased and six significantly decreased bacterial species in Lgals4-KO mice (Fig.1C), including those identified by ANCOM-BC. No significant variations in relative abundances were detected at the phylum to family level (Fig.S2B-C). Overall, these results indicate a significant alteration in gut microbiota in Lgals4-KO mice.

**Fig. 1.**
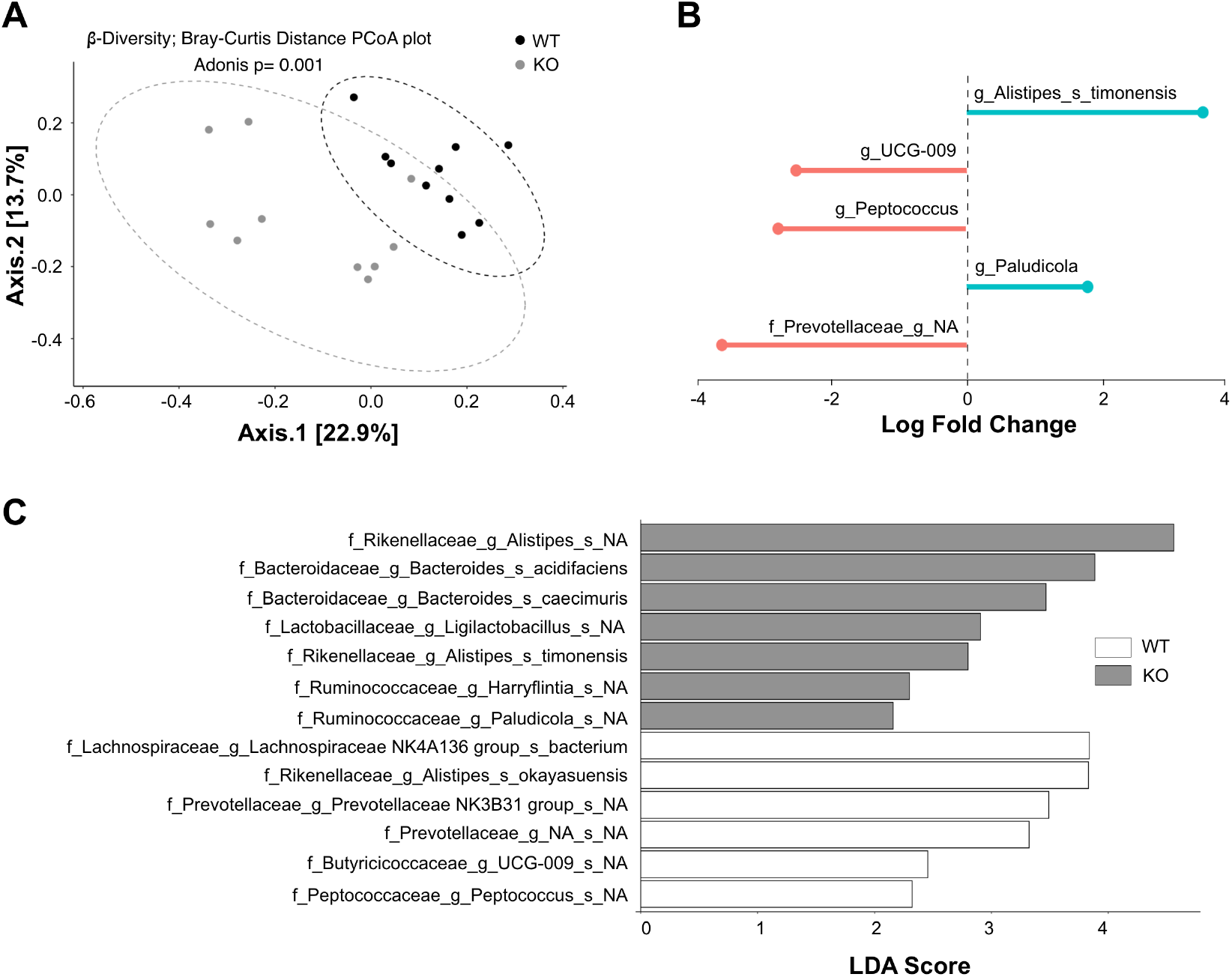
Lgals4-KO mice have an altered microbiota composition. (A) Principal coordinate analysis (PCoA) plot of bacterial beta diversity based on the Bray-Curtis distance. Gut microbiota samples from WT (black dots) and Lgals4-KO (grey dots) mice show a significantly different distribution between groups (Adonis, p=0.001). Ellipses indicate 95% confidence interval, and the percentages in parentheses are the proportion of variation explained by the PCoA axis. Animals used: WT n=10, KO n=10. (B) Lollipop plot shows the differentially abundant taxa identified by ANCOM-BC analysis. Each line indicates a differentially abundant taxon, with the line length reflecting the magnitude of the log fold change and a circle at the end marking the exact value. Lgals4-KO enriched and depleted taxa are represented as blue and red lines, respectively. (C) Differentially abundant taxa identified by LEfSe analysis. Histogram representation of the LDA scores for the differentially abundant taxa between WT (white bars) and Lgals4-KO (grey bars) groups. The length of the bar represents the log10 transformed LDA score.

### Memory formation deficits in Lgals4-KO mice

Changes in the gut microbiota composition can influence brain function as seen in microbiota-depleted models (34). Therefore, we evaluated whether cognitive processes (memory) and mood-related behaviors were altered in Lgals4-KO mice. Lgals4-KO mice subjected to the Y-maze Forced Alternation test (Fig.2A) spent significantly less time exploring the novel arm compared to WT mice (Mann-Whitney U test, WT, p=0.0002) (Figs.2B-C). Additionally, Lgals4-KO mice took a significantly longer time to enter the novel arm for the first time (Mann-Whitney U test, p=0.0439) (Fig.2D). While WT mice chose the novel arm for their first entry 82.0% of time, Lgals4-KO mice did so only 64.9% of the time. However, this difference did not reach statistical significance (Chi-square test, p=0.0691) (Fig.2E). These results suggest an impaired memory formation in Lgals4-KO mice. Similarly, in the Novel Object Recognition (NOR) test, Lgals4-KO mice spent less time exploring the novel object, with novelty preference scores of 61.5% and 46.3% for WT and KO, respectively (Unpaired t-test, p=0.0100) (Fig.2F-H), corroborating an impaired memory formation. Lastly, in the Object Location test, where WT mice are expected to explore the novel location longer, our control strain unexpectedly showed no preference, making this test inconclusive (Fig.S3A).

**Fig. 2.**
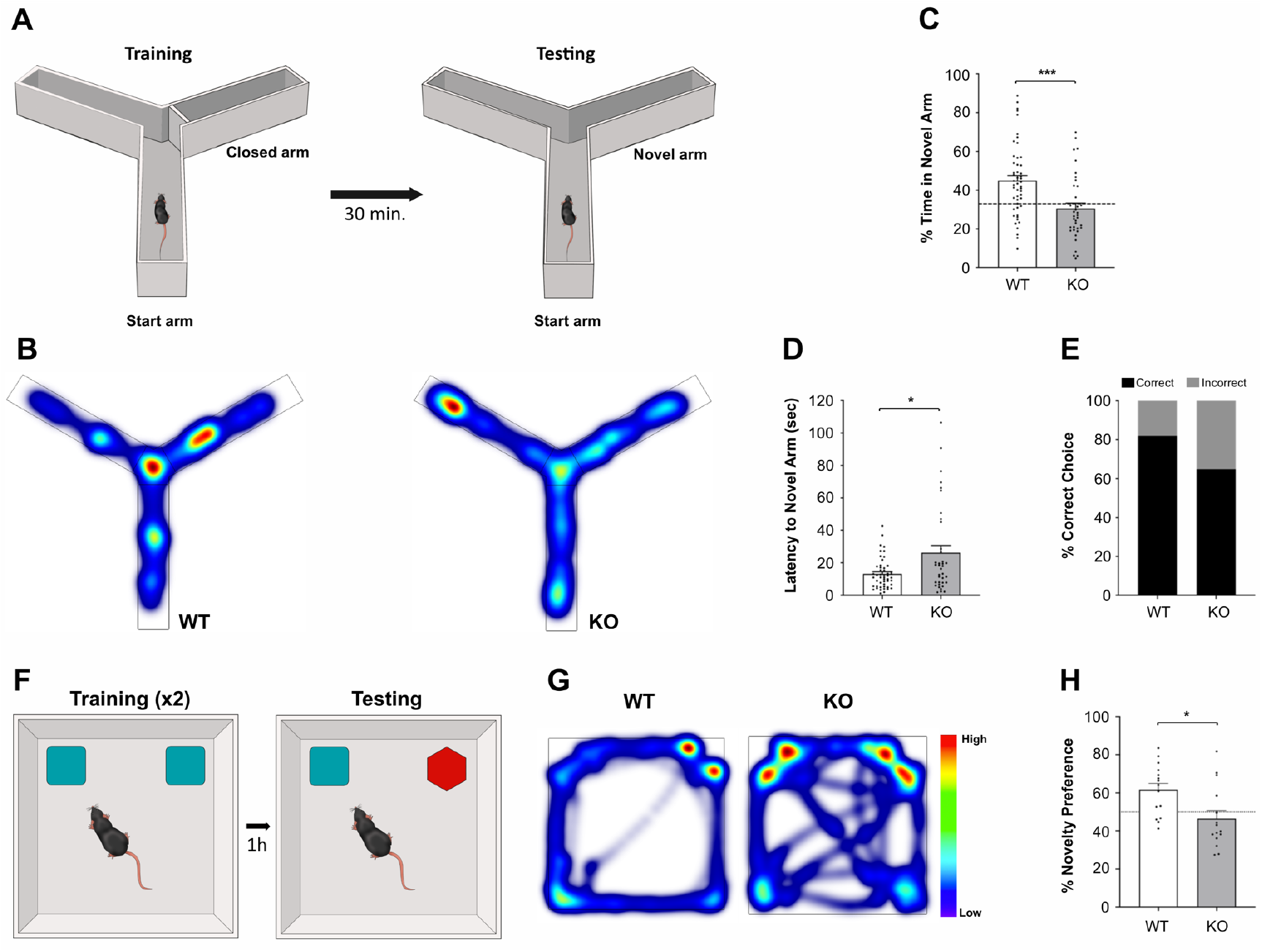
Lgals4-KO mice display an impaired hippocampus-dependent spatial memory formation. (A) Schematic representation of the Y-maze test. (B) Representative heatmaps of the cumulative time mice spent exploring each of the three arms of the Y-maze for WT (left) and KO (right) mice during the testing session. Warm and cold colors are indicative of more and less time spent exploring each arm, respectively. (C) Bar graph represents the percentage of total time that mice spent exploring the novel arm. Lgals4-KO mice spent significantly less time exploring the novel arm (Mann-Whitney U test; ***p=0.0002). The dashed line at 33% represents the percentage of time mice would randomly explore any of the three arms. (D) Bar graph shows that Lgals4-KO mice display a higher latency to explore the novel arm (Mann-Whitney U test; *p=0.0439). (E) Stacked bar graph represents the percentage of correct and incorrect first arm entry choices. There is no difference in the proportion of correct first arm entry choices between WT and Lgals4-KO mice (Chi-square test; p=0.0691). (F) Schematic representation of the NOR test. (G) Representative heatmaps of the cumulative time mice spent exploring the novel and familiar objects for WT (left) and KO (right) mice. Warm and cold colors are indicative of more and less time spent exploring each object, respectively. (H) Bar graph showing that Lgals4-KO mice spent significantly less time exploring the novel object expressed as a percentage of Novelty Preference (Unpaired t-test; *p=0.0100). The dashed line at 50% represents the percentage of time mice would randomly explore any of the two objects. All represented data are means + SEM. Animals used for the Y-maze test: WT n=50, KO n=37; for NOR test: WT n=15, KO n=15.

Depression-like behavior was examined using the Porsolt forced swim and the tail suspension tests. In both tests, WT and Lgals4-KO mice spent similar percentages of time immobile (Fig.S3B-C). Finally, we used the elevated zero maze test to assess anxiety-like behavior. As before, both strains spent comparable time in the open zones (Fig.S3D).

Taken together, our results show that Lgals4-KO mice present an impaired recognition and spatial working memory, with no evident alterations in anxiety- or depressive-like behaviors.

### Impaired LTP induction in the hippocampus of Lgals4-KO mice

To test whether memory deficiencies in Lgals4-KO mice were associated with altered LTP, we performed extracellular recordings of excitatory postsynaptic field potentials (fEPSP) in acute brain slices. The fEPSPs were recorded in the CA1 region of the hippocampus while Schaffer collaterals were stimulated to elicit synaptic NMDAR-dependent LTP responses (Fig.3A). The response upon LTP stimulation was significantly greater in control compared to knockout slices (Mann-Whitney U test, p=0.0006) (Fig.3B-C), while basal synaptic function was normal according to the unaltered basal transmission in Lgals4-KO (Fig.S4).

**Fig. 3.**
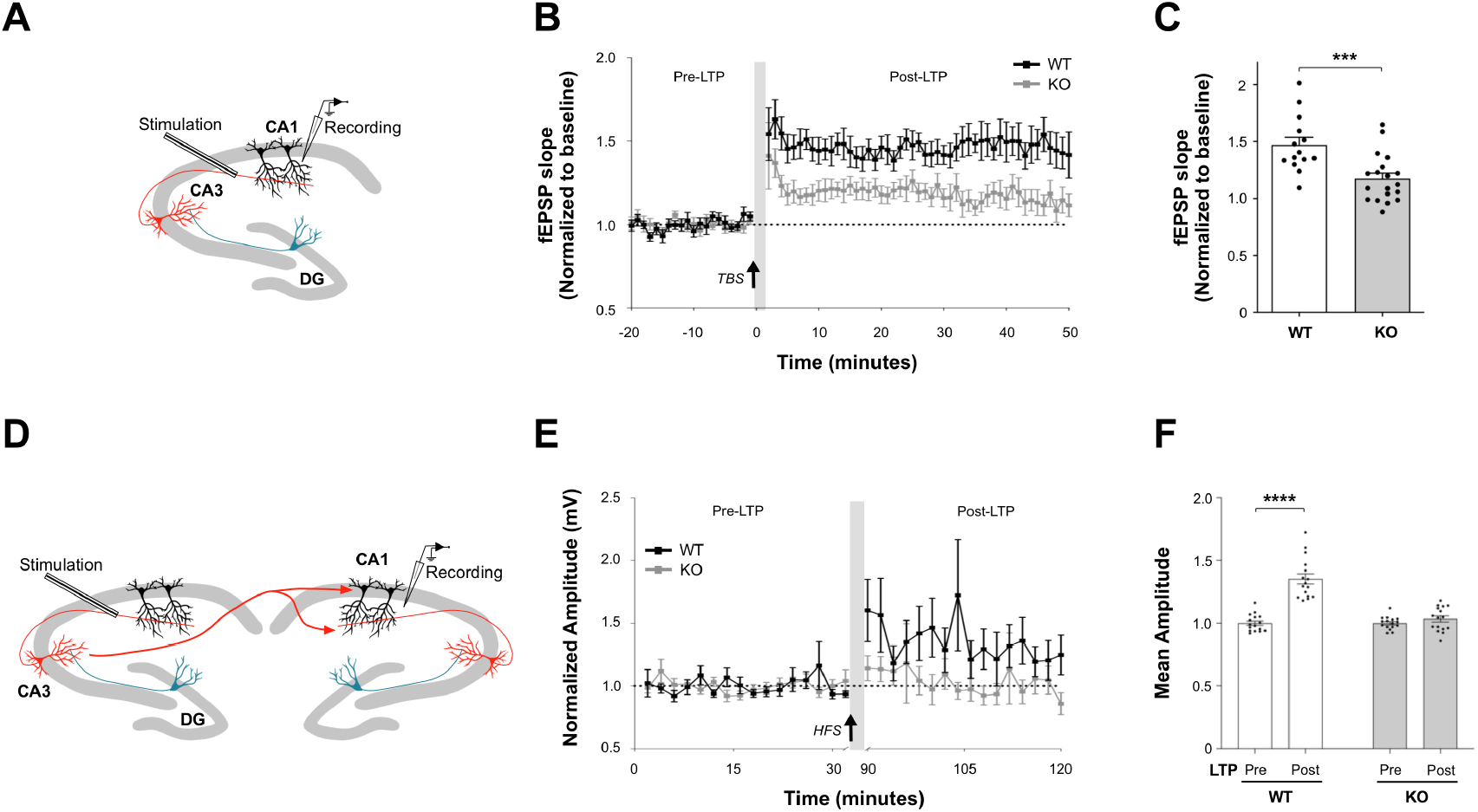
Impaired long-term potentiation of synaptic strength in Lgals4-KO mice. (A) Schematic representation of the induction and recording of LTP in acute hippocampal slices. Responses were evoked by stimulating Schaffer collaterals and recorded from CA1 *stratum radiatum*. (B) Time course of LTP induced by theta-burst stimulation (TBS, black arrow) in WT and Lgals4-KO hippocampal slices. (C) Bar graph shows a statistically significant reduction in the average change in fEPSP slope from the last 25 min of the recording in Lgals4-KO slices compared to control (Mann-Whitney U-test, ***p=0.0006). Dots represent the number of recordings (WT n=14 from 14 slices; KO n=19 from 17 slices) obtained from 9 mice per condition. (D) Diagram of a mouse hippocampus with electrodes in place for the induction and recording of LTP in anesthetized animals *in vivo*. Stimulation was performed in CA3 and responses were recorded in the contralateral CA1 region. (E) Time course of LTP induced by high-frequency stimulation (HFS, black arrow) in WT and Lgals4-KO mice. Pre- and post-LTP responses were recorded with a 60 min gap after stimulation. (F) Bar graph shows a statistically significant increment in the amplitude of post-LTP responses in WT mice but not in Lgals4-KO mice (Wilcoxon signed rank test, WT ****p<0.0001; Paired t-test, KO p=0.2271). Dots represent the mean normalized amplitude values for each of the 16 stimuli before and after LTP induction obtained from 12 WT and 21 Lgals4-KO mice. Data are means ± SEM.

Electrophysiological responses to LTP stimulation were also measured *in vivo* by stimulating the CA3 hippocampal region of anesthetized mice and recording at the pyramidal layer of contralateral CA1 (Fig.3D). Consistent with *ex vivo* results, control mice produced a significantly greater response upon LTP stimulation compared to normalized pre-stimulation values (Mann-Whitney U test, p<0.0001), while in knockout mice the response was indistinguishable from basal level (Fig.3E-F). Together, these electrophysiological results demonstrate that Lgals4-KO mice present alterations in the CA3-CA1 hippocampal pathway that significantly reduce the capacity of LTP generation.

### WT and Lgals4-KO mice have comparable hippocampal neuronal densities and myelin expression and distribution

As a first approach to explaining the impaired memory phenotype and the deficient LTP induction in Lgals4-KO mice, we examined potential morphological or cellular alterations in the Lgals4-KO hippocampus. However, WT and Lgals4-KO animals presented comparable hippocampal volumes (Fig.S5B-D) and similar cell densities of neurons (Fig.S5A, S5C-D), oligodendrocytes (Fig.S6B), astrocytes and microglia (Fig.S10). Given that myelin plasticity can be altered by gut microbiota changes (35–37) and may, in turn, affect memory formation by altering electrophysiological properties (38), we investigated whether myelin remodeling contributes to the observed phenotype. In our model, no differences were found in the quantity or distribution of Lgals4-KO hippocampal myelin using Black Gold staining (Fig.S6A). Consistently, immunohistochemical staining of major myelin markers, myelin basic protein (MBP) and proteolipid protein 1 (PLP) (Fig.S6C), showed no differences in expression or distribution between strains. RNA-sequencing of hippocampal tissue revealed that only 5 genes annotated as myelin-related, present minimal differential expression between strains (Fig.S6D; Supplemental RNAseq data). Consistently, the hippocampal expression of relevant myelination markers measured by RT-PCR (Fig.S6E) or by western blot were also similar (Fig.S6F).

In all, the deficiencies in memory and LTP in Lgals4-KO mice are not attributable to any perturbation of hippocampal morphology, cellular populations, or myelin composition and distribution.

### Reduced activation of GluA1 subunit and CaMKII upon chemical LTP stimulation of Lgals4-KO neurons

The phosphorylation of the GluA1 subunit of the AMPA receptor and its delivery to the neuronal plasma membrane are main biochemical features of LTP (39,40). Upon cLTP induction, the pGluA1/total GluA1 ratio was significantly lower in hippocampal neuron membranes of Lgals4-KO mice than in WT mice (Unpaired t-test, p=0.0012) (Fig.4A). Notably, the addition of recombinant Gal-4 did not rescue this effect (Fig.4A). Consistent with these findings, immunocytochemical staining of GluA1 extracellular epitopes before permeabilization and of intracellular epitopes after permeabilization showed that the surface GluA1/total GluA1 remained unchange in Lgals4-KO neurons upon cLTP induction but significantly increased in WT neurons (Unpaired t-test, p=0.0289) (Fig.4B).

**Fig. 4.**
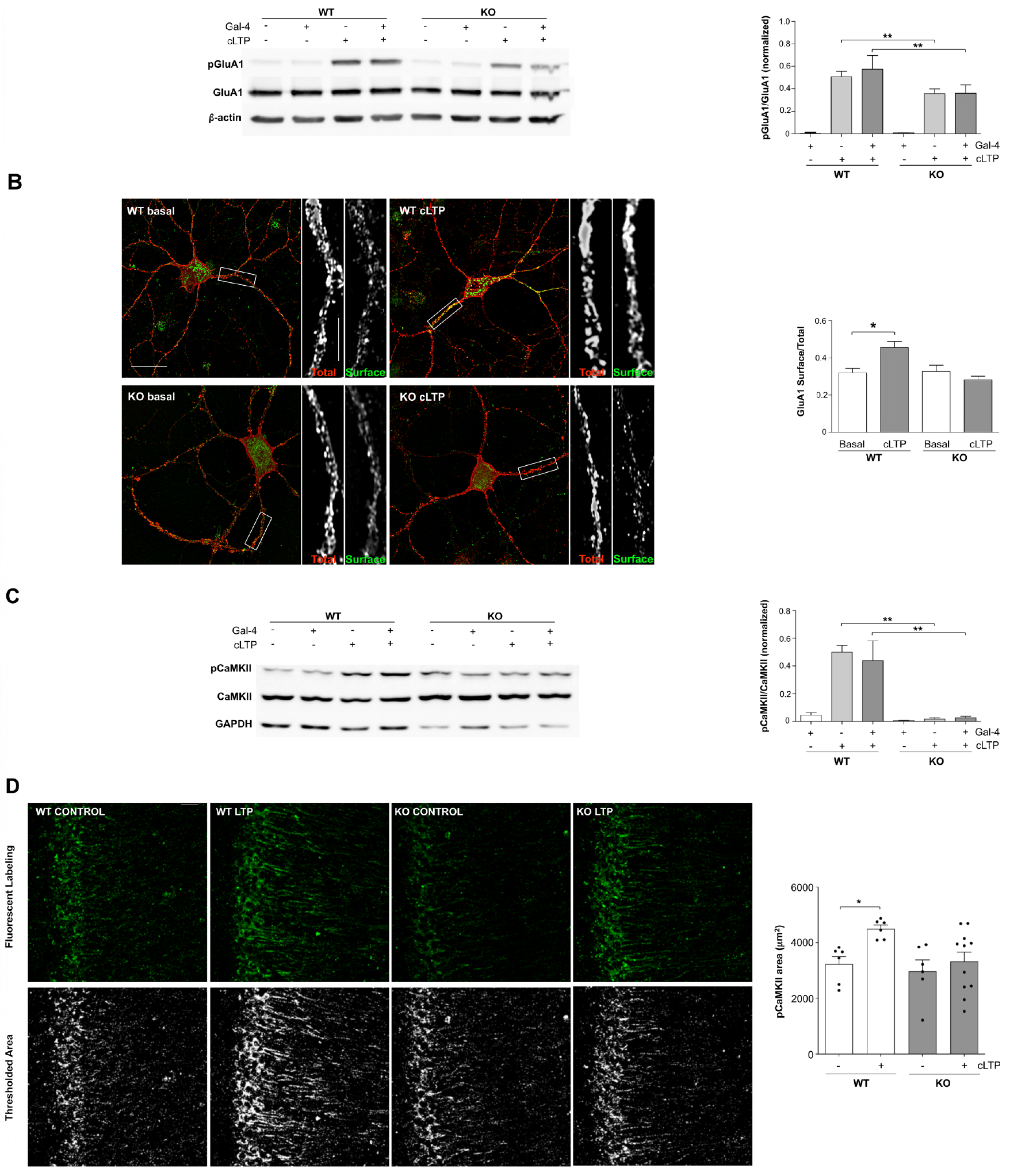
Reduced activation of GluA1 receptors and αCaMKII after LTP induction in Lgals4-KO mice. (A) Representative western blot of GluA1 S845 phosphorylation (pGluA1) before (−) and after (+) cLTP induction in WT and Lgals4-KO hippocampal neuron lysates, with or without recombinant Gal-4 treatment. Total GluA1 and β-actin are shown as loading controls. Graph shows pGluA1/GluA1 ratio in all conditions. Lgals4-KO neurons display reduced pGluA1/GluA1 levels after cLTP induction compared to WT (Paired t-test; **p=0.0012), which are not rescued by recombinant Gal-4 (Paired t-test; **p=0.0012). Bars represent the mean ratio + S.E.M. normalized to the basal conditions (Gal-4 -; cLTP -). Results were obtained from at least 3 different cultures. (B) Representative images of DIV14 cultured neurons before and after cLTP induction immunostained for surface GluA1 receptors (green, non-permeabilized) and total GluaA1 (red, after permeabilization). Magnified grayscale insets (right) correspond to dashed white rectangles. Scale bars: 25 μm (main), 5 μm (insets). Graph shows a significant increase in surface/total GluA1 ratio in WT (Unpaired t-test; *p=0.0289) but not in Lgals4-KO (Unpaired t-test; p=0.3212) cultures after cLTP induction. Bars represent the mean ratio + S.E.M. obtained from at least 3 different cultures. (C) Representative western blot of T286 phosphorylated αCaMKII (pCaMKII) before (−) and after (+) cLTP induction in WT and Lgals4-KO hippocampal neurons, with or without recombinant Gal-4. Total αCaMKII and GAPDH are shown as loading controls. Graph shows pCaMKII/total αCaMKII ratio in all conditions. Lgals4-KO neurons display reduced phosphorylated/total αCaMKII levels after cLTP induction compared to WT neurons (Paired t-test; **p=0.0084), which are not reversed by the addition of recombinant Gal-4. Bars represent the mean ratio + S.E.M. normalized to the basal conditions (Gal-4 -; cLTP -). Results were obtained from at least 3 different cultures. (D) Representative immunofluorescence images of T286-autophosphorylated αCaMKII (green) in the dorsal CA1 region of control and LTP-induced WT and Lgals4-KO mice (upper panels). Lower panels show thresholded immunopositive areas. Scale bars: 25 μm. Graph shows pCaMKII immunopositive area in all groups. LTP induction increases T286-autophosphorylation in WT but not in Lgals4-KO mice (Two-way ANOVA; effect of strain: F(1,25) = 4.291, p=0.0488; LTP induction effect: F(1,25) = 5.465, p=0.0277; Sidak’s multiple comparisons test: WT vs. WT+LTP, *p=0.0442; KO vs. KO+LTP, p=0.6959). Dots represent the mean value of all sections analysed per animal (WT n=6, WT+LTP n=6, KO n=6, KO+LTP n=11). Data are mean + SEM.

Activation of CaMKII in neurons is a central signaling step for the long-lasting consolidation of LTP downstream of AMPA receptor activation (41,42). In vitro testing of CaMKII activation showed that the normalized pCaMKII/CaMKII ratio increased after cLTP induction in cultured WT neurons (Paired t-test, p=0.0084), whereas no effect was detected in Lgals4-KO neurons (Fig.4C). As with GluA1 phosphorylation, the addition of recombinant Gal-4 failed to restore CaMKII activation in Lgals4-KO neurons (Fig.4C). Consistent with these *in vitro* findings, immunofluorescence labelling of phosphorylated CaMKII in brain slices from mice that underwent *in vivo* LTP stimulation revealed a significant increase in the CA1 hippocampal region of WT mice, while it remained unaltered in Lgals4-KO mice (Two-way ANOVA, Sidak’s multiple comparisons test: WT vs. WT+LTP, p=0.0442; KO vs. KO+LTP, p=0.6959) (Fig.4D). In all, both biochemical and immunofluorescence results indicate a deficient activation of CaMKII upon LTP stimulation in Lgals4-KO mice, consistent with the reduced AMPA receptor activation, and with the impaired LTP formation *ex vivo* and *in vivo*.

### Reduced dendritic spine density and shorter dendritic spine length in apical and basal dendrites of Lgals4-KO neurons

LTP induction and consolidation involve the functional strengthening of existing synapses and/or the formation of new ones (42–44). We hypothesized that the deficient LTP induction in the Lgals4-KO hippocampus could be caused by synaptic alterations. To investigate this, RNA sequencing analysis of hippocampal tissue was performed, and genes were classified as significantly differentially expressed if they met the criteria of P_adj_ < 0.05 and an absolute log_2_ fold change ≥ 0.1. Based on these criteria, 281 genes showed significant differential expression between mouse strains. Out of these, 51 genes were associated with synaptic functions, including 28 genes related to postsynaptic processes (Fig.S8B) (Supplemental RNAseq data). At the protein level, only trends in differential expression could be detected (Fig.S8C). Despite this, the predominant changes in postsynaptic-related genes prompted us to explore possible alterations in postsynaptic structures.

The number and structural features of dendritic spines were compared in Golgi-Cox-stained brain slices (Fig.5A-B). Spine density in both basal and apical dendrites of CA1 hippocampal neurons was quantified by counting the number of spines in length-matched dendritic segments (10 μm) (Fig.5C). Lgals4-KO neurons showed a significantly reduced spine density in both apical and basal dendrites compared to WT neurons (Apical: Unpaired t-test, p< 0.0001; Basal: Mann-Whitney U test, p< 0.0001) (Fig.5C), regardless of the sex of the mice (Fig.S9). To determine potential morphological alterations in dendritic spines, spine lengths and spine head widths were manually traced and measured in each dendritic segment. Dendritic spines from both basal and apical dendrites of Lgals4-KO neurons were significantly shorter than those of WT neurons, as shown by the relative frequency distribution of spine lengths (Kolmogorov-Smirnov test, p<0.0001) as well as by the comparison of the mean ranks of all measured spine lengths (Mann-Whitney U test, p<0.0001) (Fig.5D). In contrast, no significant differences were detected in spine head widths (Fig.5E).

**Fig. 5.**
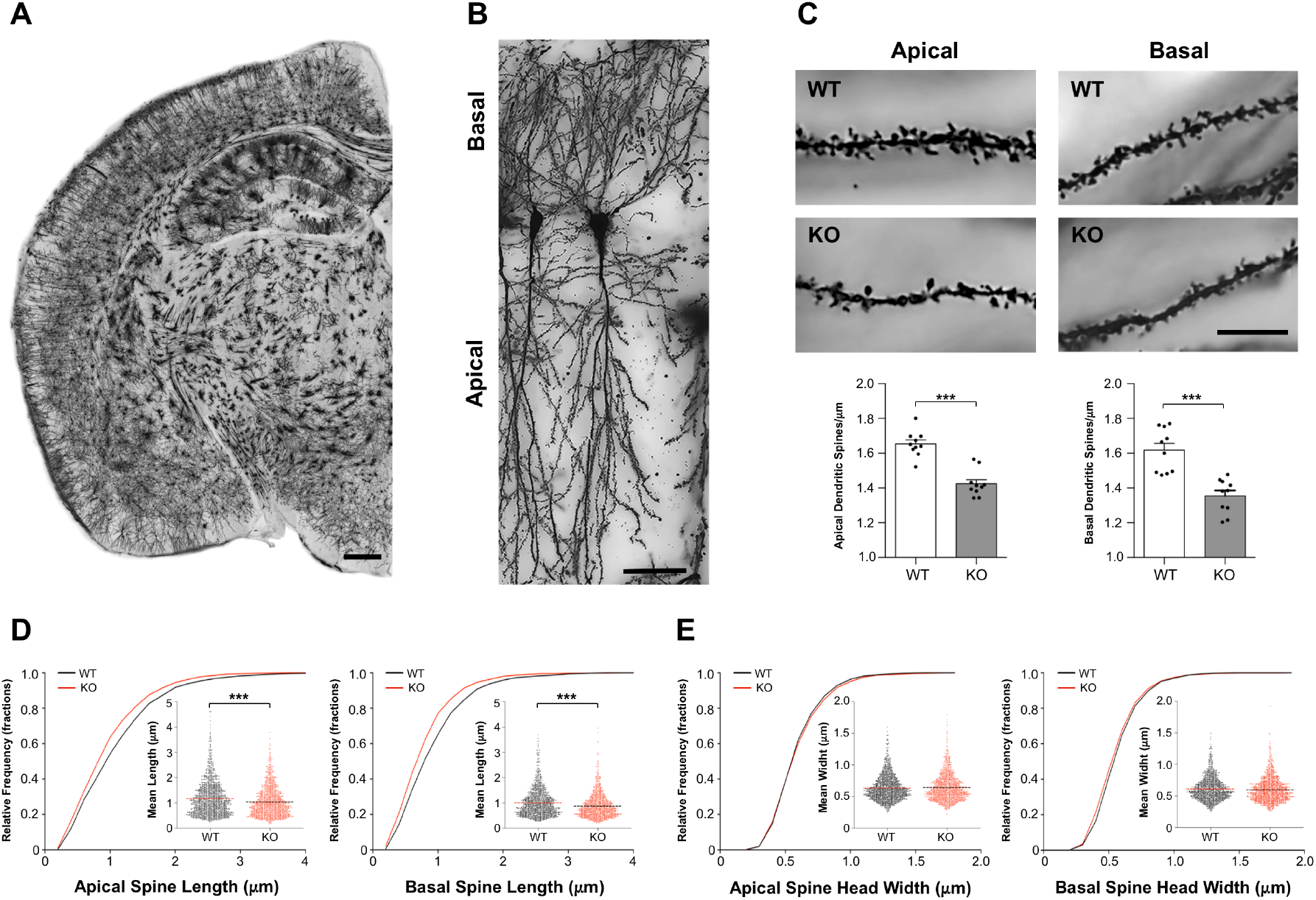
Reduced dendritic spine density and altered morphology in Lgals4-KO mice. (A) Representative mosaic image of a Golgi-Cox-stained mouse brain hemisphere. The Golgi-Cox staining technique enabled the visualization of the distribution of isolated pyramidal neurons in the hippocampal CA1 region. Scale bar: 500 μm. (B) Representative microphotograph of a fully stained CA1 pyramidal neuron showcasing its abundant arborization. The basal branches extend towards the *stratum oriens* and the apical branches reach the *stratum radiatum*. Scale bar: 50 μm. (C) Illustrative images highlighting Golgi-Cox stained apical (left panels) and basal (right panels) dendrites at high magnification. The images display numerous basal and apical spines in WT mice, while a reduced number of both apical and basal spines is evident in Lgals4-KO mice. Scale bar: 10 μm. The bottom bar graphs show the dendritic spine density within the CA1 subfield. The apical and basal dendritic spines exhibited a significantly lower density in Lgals4-KO compared to WT mice (Apical: Unpaired t-test, ***p< 0.0001; Basal: Mann-Whitney U test, ***p< 0.0001). All measured data are expressed as means + SEM of 10 animals per strain. (D) Cumulative frequency distribution of spine lengths of both apical and basal dendrites. Note that the cumulative frequency plot for spine lengths is always shifted leftward in Lgals4-KO mice, indicating a greater proportion of shorter spines (Kolmogorov-Smirnov test; *p<0.0001). The inset represents a scatter plot of each spine length value included in the analysis for both strains (Mann-Whitney U test; p<0.0001). (E) Cumulative frequency distribution of spine head widths from apical and basal dendrites. Golgi-stained spines exhibited comparable head widths across both experimental groups in both apical and basal dendrites (Kolmogorov-Smirnov test). The inset displays a scatter plot of the individual head width values measured for both WT and Lgals4-KO mice (Mann-Whitney U test).

### Increased perforated synapses and enlarged postsynaptic density areas in Lgals4-KO neurons

A reduced hippocampal dendritic spine density could be associated with fewer synapses (45), which could account for the impaired memory formation and deficient LTP in Lgals4-KO mice. We quantified synaptic density using transmission electron microscopy (TEM) images of hippocampal tissue (Fig.6A). WT and Lgals4-KO mice showed similar densities of total synapses (Fig.6A top) and of “on shaft” synapses (Fig.6A middle). In contrast, the density of high-efficiency perforated-type synapses was significantly increased in the hippocampus of Lgals4-KO mice (Unpaired t-test, p=0.0011) (Fig.6A bottom).

**Fig. 6.**
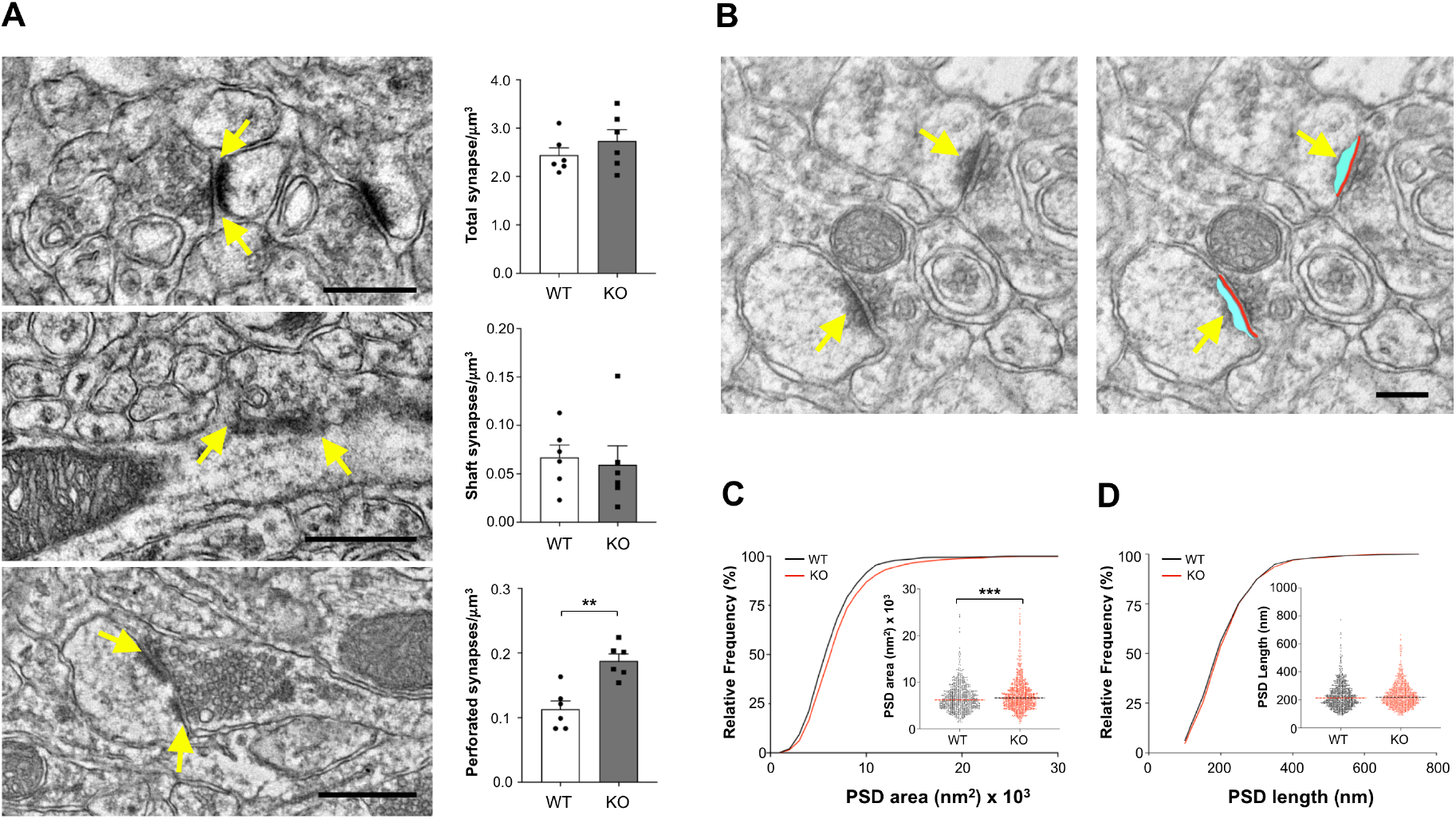
Increased number of perforated synapses and enlarged postsynaptic densities in Lgals4-KO mice. (A) Representative electron micrographs and synaptic density quantifications of total synapses (upper panels), shaft synapses (middle panels), and perforated synapses (bottom panels) in the *stratum radiatum* of WT and Lgals4-KO mice. The arrows indicate synapses marked by the presence of PSDs. Total and shaft synaptic densities were comparable between WT and Lgals4-KO mice (Unpaired t-test; p=0.3171 for total synapses, p=0.7532 for shaft synapses), while the density of perforated synapses was significantly increased in Lgals4-KO mice (Unpaired t-test, **p=0.0011). A total of 2143 synapses from 6 WT mice and 2540 synapses from 6 KO mice were analysed. Data in bar graphs are mean + SEM. Dots in bar graphs represent the mean value per animal. Scale bars: 500 nm. (B) Illustrative electron micrographs highlighting the PSD area (cyan) and PSD length (red) measured and quantified in C and D, respectively. Scale bar: 200 nm. (C) Cumulative frequency distribution of PSD areas. Note that the cumulative frequency plot is shifted rightward in Lgals4-KO mice, indicating a greater proportion of PSDs with an increased size (Kolmogorov-Smirnov test; **p=0.0026). The inset shows a dot plot of each PSD area measured in the analysis for both strains (Mann-Whitney U test; ***p<0.0001). (D) Cumulative frequency distribution of PSD lengths. The cumulative frequency plot exhibits comparable PSD lengths across both experimental groups (Kolmogorov-Smirnov test; p=0.1951). The inset displays a dot plot of each PSD length measured in the analysis for both WT and Lgals4-KO mice (Mann-Whitney U test; p=0.1045). A total of 870 synapses from 6 WT mice and 930 synapses from 6 KO mice were analysed.

In addition to synapse density, the structural features of the postsynaptic densities (PSD) are related to synaptic function (46–48). To assess postsynaptic structural differences, the cumulative frequency distributions of PSD areas and lengths were calculated from measurements obtained from TEM images (Fig.6B). Lgals4-KO mice showed a significantly larger proportion of PSDs with increased area, indicated by the rightward shift in the cumulative frequency plot (Kolmogorov-Smirnov test; p=0.0026) and by comparison of all measured PSD areas (Mann-Whitney U test; p<0.0001) (Fig.6C). In contrast, no differences were observed in PSD lengths, either in the cumulative distribution (Kolmogorov-Smirnov test; p=0.1951) or in the comparison of mean ranks (Mann-Whitney U test; p=0.1045) (Fig.6D).

## DISCUSSION

Microbiota-depleted models, such as germ-free or antibiotic-treated mice, have been used to establish microbiota influence in brain physiopathology (2,12,13,34). However, these models fail to reflect physiological conditions (49,50), where gradual changes in endogenous microbiota, occurring throughout an organism’s lifespan, could play a relevant role in brain function regulation.

The intestine exhibits the highest expression of Gal-4, where it controls infections by pathogenic self-antigen-bearing bacteria (23,26). We show distinct differences in bacterial populations between WT and Lgals4-KO mice, revealing for the first time a role for Gal-4 also in regulating endogenous microbiota. This finding prompted our hypothesis that Lgals4-KO model could provide physiologically relevant insights on the interplay between microbiota and brain function.

Lgals4-KO mice exhibited impaired performance in spatial memory formation and learning tests suggesting a connection between Gal-4 function and hippocampal-dependent cognitive processes. Notably, no alterations in mood-related behaviors (depression or anxiety) were observed across the behavioral assays applied. While memory formation can be affected by altered myelination (51,52), we demonstrate that myelin integrity in the Lgals4-KO hippocampus remains unchanged, consistent with our previous observations in the brain cortex (53), thereby excluding myelin abnormalities as a contributing factor.

A central mechanism underlying memory formation is hippocampal LTP (54). LTP induction in Lgals4-KO mice resulted in significantly reduced responses both in brain slices and in living animals. Since LTP depends on (and is proportional to) the retention of AMPA receptors at the membrane, which is regulated by the phosphorylation of their GluA1 subunit (39,40,55–57), the lower increase in pGluA1 after cLTP induction, along with the reduced amount of the receptor on the neuronal membrane in Lgals4-KO neurons, likely accounts for the deficient synaptic potentiation observed. Furthermore, Lgals4-KO neurons showed reduced CaMKII activation upon cLTP induction, as corroborated by decreased pCaMKII signal in immunofluorescence staining of brain slices obtained from LTP-stimulated animals. Given that AMPA receptor-stimulated CaMKII activation and downstream signaling pathways (i.e. RAS-ERK) are involved in memory formation (41,58,59), our results are consistent with the observed neurological phenotype of memory deficit and impaired LTP.

Considering that LTP is a synapse-dependent process, we sought to identify synaptic alterations that could underlie LTP failure. Transcriptomic analyses revealed significant differential expression for 51 synaptic-related genes in the hippocampus, of which up to 28 were postsynaptic. However, these differences were modest and compatible with an indirect effect in the hippocampus likely originating from the lack of Gal-4 in the intestine, as we hypothesized. The inability of recombinant Gal-4 to rescue WT levels of activated AMPA receptor or CaMKII upon cLTP induction, combined with the very low (if any) expression of Gal-4 in WT brain according to both the literature and to our own results (Supplemental Information), reinforce the notion of an indirect effect.

In parallel with the postsynaptic gene expression changes, Lgals4-KO CA1 neurons exhibited reduced dendritic spine length and density compared to controls. These effects are unlikely to result from increased microglial-driven spine pruning, as microglia is similar in both strains. In such a low dendritic spine density background, compensatory mechanisms such as increased in synaptic efficiency could explain the normal basal neuronal activity shown by electrophysiological measurements in Lgals4-KO mice. Although we cannot determine whether the morphological differences are a primary consequence or an adaptive response, spine neck shortening (60), increased presence of high-efficiency perforated synapses (61), and larger PSD areas (46–48) collectively contribute to maintaining basal synaptic activity. However, these adaptations may constrain further synaptic potentiation in response to strong physiological demands, such as LTP stimulation, resulting in the cognitive deficits seen in Lgals4-KO mice.

The absence of Gal-4 led to significant shifts in bacterial genera and species previously associated with different neurological conditions. Specifically, reduced levels of the genus *Peptococcus* are associated with sleep disorders (62), depression (63), and memory deficits (64,65). Additionally, *Paludicola* and *Butyricicoccaceae_UCG009* genera known to be altered by a high fat/high sugar diets (66), were respectively increased and decreased in Lgals4-KO mice fed a standard diet, implicating these taxa in synaptic deficits synaptic deficits and memory dysfunction. Low levels of *Bacteroides acidifaciens* and *Bacteroides caecimuris* are related to cognitive decline as well as to anxious and depressive behaviors in obese-diabetic mice (67,68) or after chronic stress (69,70). Increased levels of these bacterial species in Lgals4-KO mice may contribute to their normal performance in mood-related behavioral tests, although this increase would be insufficient to reverse memory deficits. Furthermore, Lgals4-KO mice present reduced *Alistipes okayasuensis*, related with poor outcomes in the DSS colitis mouse model (71). To our knowledge, this is the first report suggesting a connection between variations in this species and cognitive impairment.

The mechanisms and extent to which these intestinal microbiota alterations signal through the gut-brain communication pathways remain unanswered questions. Addressing these aspects will require metagenomic analyses of the microbiota and metabolomic studies. Nevertheless, our findings indicate that the altered profile of intestinal microbiota could underlie the observed synaptic-associated memory and cognitive deficits in Lgals4-KO mice. Moreover, the fact that this occurs in the absence of pathogenic bacterial infections, defines a new role for Gal-4 in the regulation of commensal bacteria, and suggests that precise and chronic alterations of the endogenous microbiota could contribute to the development of specific neurological pathologies. Uncovering these relationships could open novel, microbiota-targeted therapeutic strategies for neurological diseases.

## Supporting information

Brocca et al_10_April-2025 Supplemental_Info

Brocca et al_10_April_2025 Supplemental RNAseq data

## ACKNOWLEDGMENTS AND DISCLOSURES

We want to thank the staff of the microscopy, proteomics, and animal house facilities at the Hospital Nacional de Parapléjicos –SESCAM- (Toledo, Spain) for their valuable technical support, and to Dr. Rafael Luján for his helpful advice on TEM images interpretation. This work was supported by (1) grant PID2021-125428OB-I00; funded by MCIN/AEI/10.13039/501100011033 and by “ERDF A way of making Europe”; (2) grant SAF2017-83821-R; funded by MCIN/AEI/10.13039/501100011033 and by “ERDF A way of making Europe”; (3) grant PID2019-105020GB-100; funded by MCIN/AEI/10.13039/501100011033 and by “ERDF A way of making Europe”; (4) grant PID2023-149770NB-I00; funded by MICIU/AEI/10.13039/501100011033 and, as appropriate, by “ERDF/EU (5) grant SBPLY/17/180501/000250; funded by Consejería de Educación, Cultura y Deportes de Castilla La Mancha (Spain); (6) AMR holded a predoctoral fellowship grant SBPLY/19/180501/000407, funded by “Consejería de Educación, Cultura y Deportes de Castilla La Mancha”, and co-funded by the European Social Fund (Fondo Social Europeo “El FSE invierte en tu futuro”).

All study data are included in the article and in *Supplemental information* or in *Supplemental RNAseq data*.

All authors report no potential conflicts of interest.

This article has been posted as preprint on *bioRxiv*

