## Supplementary material for "Deficient Memory, Long-Term Potentiation and Hippocampal Synaptic Plasticity in Galectin-4-Deficient Mice": Brocca et al_10_April-2025 Supplemental_Info

3- Laboratory of Synaptic Plasticity Mechanisms and Contribution to Cognitive Function. Department of Molecular Neuropathology, Centro de Biología Molecular Severo Ochoa (CSIC-UAM), Madrid, Spain.

4- Genomics Unit, Centro Nacional de Investigaciones Cardiovasculares (CNIC), Melchor Fernández Almagro, 3, 28029 Madrid, Spain.

5- Grupo de Investigación Multidisciplinar en Cuidados, Facultad de Fisioterapia y Enfermería, Universidad de Castilla-La Mancha, Toledo, Spain

**This file includes:**

- Extended Experimental Procedures
- Supplementary Figures S1-S12
- Supplementary Tables S1 and S2
- References

### Extended Experimental Procedures

#### Animals

The Gal-4 knockout mice strain (C57BL/6NJ-Lgals4$^{em1(IMPC)J}$/J), was purchased from the Jackson Laboratories. Homozygous animals are viable, fertile, and show normal development and a normal life span. C57BL/6NJ mice were used as control. All mice were bred and maintained in a temperature- and humidity-controlled environment, with continuous access to food and water, under a 12 h light/dark cycle, within the officially accredited animal facilities of the Hospital Nacional de Parapléjicos-Research Center. Genotyping was performed by PCR using genomic DNA extracted from ear clip samples. Primers for Gal-4 were: WT reverse GGG ACT GTC TCA CCG TCT CA; KO reverse GCA ATC TGA ATC ACA CAG TGG, and common forward TCT CCT ATT GAC CCA TAT CCT TAG. All experimental procedures involving animals were approved by the Animal Bioethics Committee (CEBA) of the Hospital Nacional de Parapléjicos-Research Center.

#### Recombinant galectins and antibodies

Human galectin-4 (Gal-4) was obtained by recombinant production, purified by affinity chromatography on lactosylated Sepharose 4B, and controlled for purity by 1D- and 2D-gel electrophoresis and mass spectrometry, and for activity by solid-phase and cell binding assays (1,2).

For detailed information about the antibodies used in this work see Table S1.

#### Protein extracts and Western Blot

Brain hippocampus and intestine tissues from Lgals4-KO and WT mice were homogenized in glass-teflon potters in PBS supplemented with a protease inhibitor cocktail (Sigma). The homogenates were ultrasonicated and cleared by 15 min centrifugation at 13,000 rpm. Supernatants were considered as total extracts.

To obtain membrane extracts, hippocampal neuron cultures, previously submitted to cLTP induction or not, were washed with scrape solution [25 mM MES buffer, pH 7.0; 2 mM EDTA supplemented with protease inhibitor cocktail (Sigma) and phosphatase inhibitors 1 mM phenyl-methyl sulphonyl fluoride (PMSF) and 1 mM sodium ortovanadate] at 4°C and scraped from the plate surface in the same buffer using a plastic scraper. Cell suspensions were lysed by 10 up-and-down strokes through a 22-gauge needle and spun down by 10 min centrifugation at 1.5 x g. Post-nuclear supernatants were taken up to 3.5 ml of the same buffer and ultracentrifuged at $10^5$ x g for one hour at 4°C. Membrane pellets were then resuspended in PBS-0.2% SDS supplemented with phosphatase inhibitors as before.

Protein concentrations were measured by the bicinchoninic acid assay (BCA; Bio-Rad). The extracts were further analyzed by western blot (WB) using the antibodies indicated in each case, with GAPDH as loading control. HRP-conjugated anti-species antibodies were used as secondary antibodies, and chemiluminescence was used for quantitation (Plus-ECL; Perkin-Elmer).

**Immunoprecipitation and proteomic analysis**

*Immunoprecipitation and SDS-PAGE:* Tissues were lysed at 4°C in RIPA lysis buffer with a protease inhibitor cocktail. After 10 min centrifugation at 15,000 x g, the insoluble pellet was removed. The soluble fraction was incubated with the appropriate antibody for 1 h at 4°C. Immunocomplexes were adsorbed on Protein A or Protein G-Sepharose 4B (GE Healthcare, Amersham Place, UK). Samples of the flow-through and washes were reserved for analysis. Bound complexes were eluted with Laemmli sample buffer.

For immunoblot analysis, samples were processed as indicated in *"Protein extracts and Western Blot"* section. For proteomics, samples were resolved by SDS-PAGE in 12% polyacrylamide gel, stained with Coomassie blue, and bands were manually excised.

*In Gel digestion:* The digestion was performed according to Schevchenko et al. (3) with minor modifications: gel slices were incubated with 10 mM dithiothreitol (Sigma Aldrich) in 50 mM ammonium bicarbonate (99% purity; Scharlau) for 30 min at 56°C. After reduction, alkylation with 55 mM iodoacetamide (Sigma Aldrich) in 50 mM ammonium bicarbonate was carried out for 20 min at room temperature (RT). Gel plugs were washed with 50 mM ammonium bicarbonate in 50% methanol (gradient, HPLC grade, Scharlau), rinsed in acetonitrile (gradient, HPLC grade, Scharlau) and dried in a Speedvac. Dry gel pieces were then embedded in Trypsin/Lys-C Mix, Mass spec. grade (Promega, Madison, WI, USA) at a final concentration of 12.5 ng/µL in 20 mM ammonium bicarbonate. After digestion at 37°C overnight, peptides were extracted with 60%

acetonitrile (ACN) in 0.1% trifluoroacetic acid (for LCMS; Scharlau) and the samples were resuspended in 4.5 µL of 98% water with 2% formic acid (FA) and 2% ACN.

*LC-MS/MS and database searching:* To perform the LC-ESI analysis, a TEMPO nano LC system (ABSciex) combined with a nano LC Autosampler coupled to a modified triple quadrupole (ABSciex 4000 QTRAP LC/MS/MS System) was used. The column used was a C18 Ion Exchange (NTCC-360/75-3-123). Injections of 4 µL containing the entire sample from the band were made per sample. Peptides were eluted at a flow rate of 300 nL/min in a gradient of phase A and phase B (98% ACN/2% water, 0.1% FA), following these steps: 5%-50% B for 70 min, 50%-95% B for 1 min, 95% B for 3 min, 95%-5% B for 2 min. Column was then regenerated with 5% B for 15 min. Both TEMPO nano LC and 4000 QTRAP system were controlled by Analyst Software v.1.5.2. The mass spectrometer operated in positive ion mode with unit resolution in both Q1 and Q3, an ion spray voltage of 2800V and a nanoflow interface heater temperature of 80°C. Source gas 1 and curtain gas were set to 30 and 20 psi, respectively and nitrogen was applied as curtain and collision gases.

*Data Analysis:* Data from MS/MS spectra were searched using MASCOT Versions 2.5 (Matrix Science Ltd., UK) against the UniProt Fasta dataset downloaded from UniProt Mus Musculus (62656 sequences; 27997905 residues). Analysis performed with maximum missed-cut value set to 1. Following features were used in all searches: (i) variable methionine oxidation, (ii) fixed cysteine carbamidomethylation, (iii) precursor mass tolerance of 0.8 Da, (iv) MS/MS tolerance of 0.8 Da, (v) significance threshold (p) below 0.05 (MudPIT scoring) and (vi) minimal peptide Mascot score of 34.

**Behavior tests**

*Y-Maze:* C57BL/6NJ control and Lgals4-KO mice were tested in a Y-Maze apparatus (Noldus). The maze consisted of three identical arms (35 x 6 x 10 cm) set at 120-degree angles from each other. The “Entry”, “Familiar” and “Novel” arms were randomly assigned for each animal tested. A visual cue was placed at the end of each arm to increase spatial recognition. Mice underwent a single 5 min training session in which each animal was placed facing the end of the entry arm with the novel arm closed. Therefore, mice were only able to freely explore the entry and familiar arms. After training, mice were returned to their cages for a 30 min inter-trial interval before the testing session began. During testing, the previously blocked novel arm was opened, and mice were allowed to freely explore all arms for 5 min. Both training and testing sessions were recorded. The videos were later analyzed by an observer blind to the genotype of the mice using the Ethovision XT software. For the first 2 min of the testing session, the number of arm entries, latencies and time spent in each arm were calculated, starting from the moment the animal left the entry arm (defined as having all four paws outside the arm). This ensured that the time spent exploring the novel and familiar arms was not influenced by variation in exit time from the entry arm (4). The 2 min analysis period corresponds to the peak of exploratory activity (5–7). Mice were considered to be in an arm only when the four paws were inside. Time spent in the novel arm was calculated as a proportion of the time spent in the novel arm relative to total time spent in all arms during the 2 min analysis period. The following exclusion criteria were applied: 1) mice that remained immobile and did not leave the entry arm in the first 1 min of the testing session, and 2) mice that did not enter each arm at least once during the 2 min period analyzed in testing session.

*Novel object recognition task (NOR) and object location task (OL):* The NOR and OL tests assess memory based on the spontaneous tendency of mice to explore a novel object or location over a familiar one. The testing arena consisted of a square open field (50 x 50 x 50 cm, Noldus, Wageningen, The Netherlands) with visual cues on the walls. Mouse movement was recorded and tracked using the EthoVision XT detection system (Noldus, Wageningen, The Netherlands). The experimental workflow spanned three consecutive days: the open field test which was used as habituation phase, followed by the novel object recognition test and, finally, the object location test. On day one, mice were allowed to freely explore the arena for 15 min. The total time spent in each quadrant, in the center and along the border of the arena, and the movement time were calculated as control parameters to exclude possible environmental or locomotor biases. On day two, the NOR test was performed. Two objects with similar dimensions and shape, but different materials (a plastic bottle and a glass jar filled with aluminum foil) were used. This test consisted of two training phases and a testing phase, each lasting 10 min with a 1 h inter-trial interval. During training, mice were placed in the center of the arena with two identical objects in adjacent corners, which were randomly changed for each mouse tested to prevent positional bias. After the retention period, during the testing phase, one randomly selected object was replaced with a novel object in the original location.

The OL test was performed 24 h after the NOR test, using a different object (plastic flask with wood shavings). Like the NOR test, it included a training phase and a testing phase, each lasting 10 min with a 1 h inter-trial interval. During the training session, mice were allowed to explore two identical objects placed in fixed locations. After the retention period, one of the objects was relocated to the opposite corner. The time spent exploring each object was recorded. Exploration was defined as close contact with the object, with the nose pointed toward it and within 2 cm of its surface. Object preference was expressed as a percentage (time exploring the novel object or location divided by the total time exploring both objects). To exclude mice with inherent position biases during the training phase, the Object Bias Score (OBS) (time exploring object A divided by total time exploring both objects) was calculated. Mice with an OBS below 25% or above 75%, or those that explored both objects for less than 10 s, were excluded from further analysis. Between sessions, the arena was thoroughly cleaned with 70% ethanol to minimize olfactory cues.

For this study, mice aged 3 to 9 months were used (33 WT females, 21 KO females, 17 WT males and 16 KO males for Y-Maze test; 6 WT females, 7 KO females, 9 WT males and 8 KO males for NOR test; 6 WT females, 3 KO females, 8 WT males and 5 KO males for OL test).

*Tail Suspension Test*: The tail suspension test was performed as previously described (8) using a custom-made, three-walled gray plastic apparatus (55 cm height x 60 cm width x 11.5 cm depth). The interior of the apparatus was divided into four compartments (each 15 cm wide), allowing for the simultaneous testing of four mice. Mice were suspended from a bar positioned at the top of the apparatus using a securely adhered tape strip attached to their tails. To prevent tail climbing, plastic hollow cylinders (climb stoppers) were placed at the base of the tails. Once suspended, mice behavior was recorded for 6 min. Total immobility time, defined as the amount of time (> 2 s) in which the mouse remained motionless, was later analyzed by an observer blind to the genotype using the activity analysis feature of the Ethovision XT software.

*Forced Swim Test*: The forced swim test was performed as previously described (9). Mice were gently placed in glass cylinders (20 cm height x 16 cm diameter) filled with water (24 ± 1°C) to a depth of 12 cm, ensuring that they could not touch the bottom with their tails. Cylinders were separated by opaque screens to prevent visual contact between animals during the procedure. Mouse behavior was recorded during the 6 min test period, after which they were gently dried with paper towels and placed in warm cages (bedding temperature of 32 ± 1°C) for 10 min before being returned to their home cage. The total immobility time, defined as the amount of time (> 2 s) in which the mouse remained motionless or made only minimal movements to keep its head above water, was later analyzed by an observer blind to the genotype using the activity analysis feature of the Ethovision XT software.

*Elevated Zero Maze*: The test was conducted using a circular elevated zero maze (60 cm diameter, Ugo Basile) under dim red-light conditions. Mice were placed at the boundary between an open and closed zone, facing the inside of the closed zone. Mice were allowed to freely explore the novel environment, and their behavior was recorded during the 5 min test period. The time spent in the open zones of the maze was calculated for each animal by an observer blind to the genotype using the tracking feature of the Ethovision XT software. Mice were considered to be in an open zone only when all four paws were inside.

**Electrophysiology**

*Brain slices:* Acute hippocampal slices were obtained from 11–12-month-old male and female mice. Animals were anesthetized with isoflurane and quickly decapitated once they were unresponsive to tail and foot pinches. The brains were rapidly removed and submerged in ice-cold, $Ca^{2+}$ free dissection solution (in mM: 10 D-glucose, 4 KCl, 26 $NaHCO_3$, 233.7 sucrose, 5 $MgCl_2$, 0.001% (w/v) phenol red as pH indicator) previously saturated with carbogen (5% $CO_2$, 95% $O_2$). Coronal slices (350 µm), obtained by cutting the brain in the same solution with a vibratome (Leica, VT1200S), were transferred to carbogen-gassed aCSF (in mM, 119 NaCl, 2.5 KCl, 1 $NaH_2PO_4$, 26 $NaHCO_3$, 11 glucose, 1.2 $MgCl_2$, 2.5 $CaCl_2$, osmolarity adjusted to 290 mOsm) and left to recover for 1 h at 32°C. The slices were then transferred to an immersion chamber coupled to an Olympus BX51WI microscope and were continuously perfused with aCSF (3.5 ml/min) gassed with 5% $CO_2$ and 95% $O_2$, at 25°C during the entire recording session. *Recordings:* field excitatory postsynaptic potentials (fEPSPs) were recorded from CA1 *stratum radiatum* with glass Ag/AgCl electrodes filled with aCSF (0.5-1 MΩ) while stimulating Schaffer collateral fibers with tungsten bipolar electrodes (50 µs pulses, 20-250 µA). For LTP experiments, the stimulation intensity was adjusted to 30% of the maximum response, with baseline stimulation at 0.067 Hz (one stimulus every 15 s). LTP was induced using a theta-burst stimulation (TBS) protocol, consisting of 10 trains of bursts (4 pulses at 100 Hz with a 200 ms interval), repeated for 4 cycles with a 20 s inter-cycle interval. To block $GABA_A$ receptors, aCSF was supplemented with 100 µM picrotoxin in all experiments.

*Data acquisition and analysis:* Electrophysiological recordings were conducted using Multiclamp 700 B amplifiers and the 1440A Digidata A/D converter with pClamp software (Molecular Devices). Data analysis was performed using custom-made Excel (Microsoft) macros to quantify the fEPSP initial slope. For time-course representations, responses were binned in 60 s intervals and normalized to the average baseline response.

*Electrophysiology in vivo*: A cohort of 33 animals was used for electrophysiological experiments, of which 12 were wild type and 21 were Lgals4-KO. Mice were anesthetized with isoflurane (1.5-2% in $O_2$) and placed in a stereotaxic frame (SR-6 Narishige Scientific Instruments, Tokyo, Japan) for cranial surgery. Body temperature was maintained at 36°C using a homeothermic blanket (CIBERTEC, SL. Madrid, Spain). Lidocaine (2%) was administered subcutaneously in the cranial area as a local anesthetic before removing the skin and exposing the cranial bone. Small craniotomies were made over stereotaxic coordinates targeting the pyramidal layer of hippocampal CA1 in the right hemisphere (AP -1.5, ML 1.2, DV 1.2-1.5 mm) and the contralateral CA3 (AP -1.1, ML 0.8, DV 1.5-1.7 mm). The correct CA1 pyramidal layer location was confirmed by detecting the narrow strip of multiunit action potentials and the local field potential (LFP) response as a populational spike in response to contralateral CA3 stimulation. The location of the contralateral CA3 stimulating electrode was determined when optimal CA1 LFP responses were observed at lower stimulation intensities. Electrophysiological recordings from CA1 were obtained using a tungsten electrode (TM31C40KT, 4-MΩ impedance at 1 kHz or TM31A50KT, 5-MΩ impedance at 1 kHz: World Precision Instruments Inc., Sarasota FL, USA) in AC mode filtered 0.5-3000 Hz using a system composed of a pre-amplifier, an amplifier, and a filter (Neurolog, Digitimer Ltd. UK). Electrical stimulation in CA3 was applied using a concentric bipolar stainless-steel electrode, with 0.5 mm separation between outer electrode (330 µm diameter) and core electrode (100 µm diameter) (Microprobes for Life, Gaithersburg, MD, USA). Square electrical pulses (50 µs duration) were delivered using a Master-8 electrical stimulator (AMPI, Jerusalem, Israel) and an Iso-Flex isolator unit (AMPI). To determine the response threshold in CA1, stimulation intensities (0.1–1 mA) were applied to CA3 in increasing steps. The threshold intensity was defined as the lowest intensity at which a population spike (acute negative wave) in CA1 was evoked by 50% of the stimuli. LFP responses were recorded at the threshold intensity throughout the entire experiment, including the baseline period, high frequency stimulation (HFS) for LTP induction, and post-LTP period. LFP responses were collected every 2 min for 34 min during both baseline and post-LTP conditions. LTP was induced using a HFS protocol consisting of 10 trains of 10 stimuli at 200 Hz, with a 1 s inter-train interval. Post-LTP LFP responses were recorded 1 h after HFS, maintaining the same stimulation rate (1 stimulus/2 min) and duration (34 min) as the baseline protocol. Electrophysiological signals were digitalized at 20 kHz using a CED Power 1401 Plus A/D converter and recorded with Spike 2 software (Cambridge Electronic Devices. Ltd. UK). Offline analyses were carried out using Spike 2 to quantify the populational spike response magnitudes in CA1 following CA3 stimulation (unfiltered LFP signal) across the entire stimulation protocol. Response magnitudes were calculated as the peak-to-peak value (mV) from the initiation of the waveform to the maximum negative peak of the populational spike in CA1. All response magnitudes were normalized to the average baseline population responses, reducing inter-group variability and enabling improved intra- and inter-groups comparisons. Statistical comparisons (t-test) were performed to evaluate intra-group effects between baseline and post-LTP condition, and inter-group differences under the same condition.

**Immunocytochemistry and quantification of surface GluA1**

Neurons in culture were fixed with 4% paraformaldehyde (PFA) and 4% sucrose in PBS at RT. Non-specific binding was blocked with 0.2% gelatine and 1% bovine serum albumin (BSA) in PBS. To label surface GluA1, neurons were incubated for 2 h with an antibody against the GluA1 N-terminal extracellular domain. After membrane permeabilization with 0.1% Triton X-100, 0.2% gelatine and 1% BSA in PBS, total GluA1 was labelled by incubating cells for 1 h at RT with an antibody against the GluA1 intracellular C-terminal domain. After thorough washing with PBS, cells were incubated for 1 h with the appropriate anti-species antibodies conjugated to Alexa Fluor 555 or Alexa Fluor 647 (Life

Technologies). Coverslips were mounted onto glass slides using Permount (Life Technologies). Images were acquired using a confocal microscope (LSM800; Carl Zeiss), and quantification was performed using ImageJ software. Regions of interest (ROIs) corresponding to individual dendrites were selected on the total GluA1 channel. Thus, image quantification was blind with respect to the surface GluA1 channel. Image channels were split and thresholded separately. Channel background was measured in three signal-free ROIs per image, averaged, and subtracted from the total fluorescence signal. Fluorescence intensity was then measured for each ROI, and surface GluA1/total GluA1 ratios were calculated. A total of 10 images per condition, with 5-10 ROIs per image, were analyzed across three independent experiments.

**Histochemistry and Immunohistochemistry**

*Tissue sampling:* Mice (30 days old) were anesthetized with sodium pentobarbital before sequential transcardiac perfusion with 0.9% heparinized saline and 4% formaldehyde. Brains and intestines were collected. For immunohistochemistry and black gold staining, brain tissues were post-fixed overnight at 4°C, incubated in PBS for 24 h, and cryoprotected in Olmos solution (30% ethylene glycol, 0.6 M sucrose and 10 g PVP-40 prepared in 0.1 M phosphate buffer) at -20°C. Serial coronal sections (40 µm) were collected from bregma -0.94 to -2.54 mm (10), using a vibrating blade microtome (Leica VT1200/VT1200S). For dendritic spine analysis, whole brains from five male and five female mice (7 months old) of both strains were dissected and processed for Golgi-Cox staining.

For intestinal Gal-4 immunofluorescence, intestines were post-fixed in 4% PFA overnight at RT, embedded in paraffin, sectioned (2 µm-thick), and deparaffinized before immunofluorescence.

For pCamkII immunofluorescence, brains from mice of both strains and sexes subjected or not to electrophysiological LTP induction (11 KO and 7 WT per condition), were collected after electrophysiological recordings. Under anesthesia (1.5–2.0% isoflurane gas/oxygen), mice were perfused with 0.9% heparinized saline. Brains were removed, post-fixed in 4% PFA overnight at 4°C, cryoprotected in 30% sucrose in 1× PBS, frozen in isopentane pre-cooled with dry ice, sectioned on a cryostat (20 µm thick), and stored at −20°C.

*Immunostainings:* For fluorescent staining, sections were blocked with 0.2% Triton X-100, 2% BSA and 8% donkey serum for 1h, followed by overnight incubation at 4°C with primary antibodies (see Supplementary Table S1 for antibodies). Sections were washed with PBS, incubated for 90 min with fluorophore-conjugated secondary antibodies, counterstained with DAPI, and mounted with Vectashield mounting medium (Palex Medical). For immunochemical staining, sections were rinsed in PBS, incubated for 15 min in 3% $H_2O_2$ / 50% methanol to inactivate endogenous peroxidase, and blocked in 3.5% BSA / 0.3% Triton X-100. Following overnight incubation with primary antibodies at 4°C, sections were washed, incubated for 2 h at RT with biotinylated secondary antibodies, and visualized using the Vectastain Elite ABC Kit Peroxidase (Vector). Sections were dehydrated, cleared in xylene, and mounted with DPX (Sigma-Aldrich).

*Black Gold staining:* Myelin staining was performed using the Black Gold II Myelin staining kit (#AG105, Millipore), following the manufacturer's instructions. Vibratome serial sections were placed onto positively charged slides and dried completely at RT. Then, tissue sections were rehydrated with Milli-Q water for 2 min and transferred to pre-warmed 0.3% Black-Gold II solution at 60°C for 40 min until a complete myelin impregnation was achieved. After thorough washing and incubation in 1% sodium thiosulfate at 60°C for 3 min, slides were dehydrated and mounted with DPX (Sigma-Aldrich).

*Golgi-Cox staining:* For the study of spine density and spine morphology, brains were stained using FD Rapid GolgiStain™ Kit (FD NeuroTechnologies) following the manufacturer's instructions with minor modifications. Briefly, unfixed brains were immersed in the impregnation solution (A/B) for 2 weeks and later transferred to solution C for two additional days. Brains were frozen in isopentane at -80ºC, cut into 100 µm thick sections using a cryostat and mounted on gelatin-coated microscope slides. This thickness was optimal to preserve spines from dendritic segments used in the subsequent analysis. Finally, the staining process was completed using the D/E solutions, and sections were dehydrated with ethanol, cleared with xylene and mounted with DPX.

*Image segmentation and analysis:* For Olig2 in the hippocampus and Gal-4 in the intestine, images were acquired at 10X and 40X, respectively, on an Olympus IX83 microscope. EFI projection (Olympus cellSens software) was used to merge Z-stacks into a single focused image. For MBP, PLP immunolabelling and black gold staining quantification, bright field images were captured with the 20X objective of an Olympus BX61 microscope, using the newCAST software and quantified using ImageJ.

For all immunohistochemical analyses, ROIs encompassing hippocampus CA1, CA3, *stratum lacunosum moleculare* (Lm) and hilus were traced using ImageJ software, based on the Mouse Brain Atlas as a reference (Figure S6, upper left panel). To quantify the MBP and PLP1 immunoreactive area, as well as black gold-positive area, each ROI was selected, and a background subtraction was performed before applying a default threshold to segment the myelin positive area. The immunoreactive area was expressed as a fraction of the total area assessed. For Olig2-positive cell quantification, background subtraction and an OTSU threshold were applied to the selected ROI as before, and a particle analysis (size 10-100 $\mu m^2$ and circularity 0.5-1.0) was performed to obtain a count of positive nuclei. Olig2+ cell numbers were normalized to the total area analyzed. For pCamkII immunodensity measurement, Z-stacks (step size: 0.5 µm) were acquired on a confocal microscope (Leica SP5, Leica Microsystems, Germany). The subsequent analysis and image processing were performed using ImageJ. A smoothing filter was applied to each plane of the confocal stack before obtaining maximum projections. To remove the background from the image, background subtraction and arithmetic subtraction of the mean background density (obtained from a ROI traced in the *stratum oriens* of CA1) were performed. Finally, a "moments" threshold was applied and the immunoreactive area (limit to threshold) was measured in the *stratum radiatum* of CA1 of both hemispheres (excluding somas). Two fields per slice from both hemispheres, from at least three slices per animal, were analyzed. Torn tissue sections were excluded. Analyses were performed blind to experimental conditions.

For MBP and PLP1 immunohistochemical analyses, the experimental n used was WT=6 and KO n=6 animals. For Olig2 analysis, WT=4 and KO=4 animals were used. Both hemispheres per section and four sections per animal (collected every 320µm) were analyzed, and the mean value was calculated for each measured parameter.

Stereological analysis: For neuronal nuclei (NeuN) immunostaining, sections were collected every 320 µm. Images were acquired using an Olympus IX83 microscope with a 10X objective for the hippocampal volume estimation, and with a 63X oil-immersion objective for volume and neuron density quantification in CA1 and CA3.

Volume estimation. Cavalieri's method allows us to estimate the volume of a 3D solid based on a fraction of its parallel slices at regular intervals (11,12). In an adapted Cavalieri estimation, we determined volume by summing the area of the region measured in each slice and multiplying by the section interval, taking into account that sections exhibited 75% thickness shrinkage along the Z-axis following NeuN immunostaining. The area of the dorsal hippocampus and the pyramidal layer of dorsal CA1 and CA3 were analyzed by delineating the ROI with ImageJ software using the Mouse Brain Atlas (10) as reference. The volume was estimated as:

$$V = \sum A_i \cdot T \cdot \frac{1}{ssf}$$

Where: $A_i$ is the area of the region of interest in the *i-th* section of an animal, *T* is the section thickness (40 µm) and ssf is the section sampling fraction (ssf=1/8).

Neuron density (optical fractionator method). The optical fractionator method is used to estimate the total number of cells from thick sections sampled across the entire structure. This method combines the fractionator sampling design (based on systematic uniform random sampling of a known fraction of the region), with the optical disector, a 3D probe used for counting neurons (11,12). Following the protocol described in (13) for standard microscopy equipment, the number of neurons was determined in the same regions delineated for volume estimation. The counting parameters were: grid size, (120.75 x 120.75 µm), counting frame (27 x 27 µm), optical disector height (8 µm) and guard zone (1 µm). To obtain the number of neurons (*N*) in the pyramidal layer of CA1 and CA3 of each animal, NeuN(+) cells were counted in 5% of the area, and the total was estimated as:

$$N = \sum Q^{-} \cdot \frac{t}{h} \cdot \frac{1}{asf} \cdot \frac{1}{ssf}$$

Where: $\Sigma Q^{-}$ is the total number of NeuN(+) cells counted in each animal, *t* is the section thickness (40 µm), *h* is the disector height adjusted for shrinkage (32 µm), asf is the area sampling fraction (0.05) and ssf is the section sampling fraction (1/8). We also calculated the coefficient of error (CE), as described elsewhere (14), to indicate the degree of precision. A CE (m=1) below 0.08 is considered acceptable. The calculated CE for each animal was below 0.05 (range in CA1: 0.029 - 0.045; range in CA3: 0.028 - 0.040). Neuron density in each animal was calculated as the total number of neurons divided by the estimated volume of the region.

Dendritic spine analysis. Z-stack images of Golgi-Cox stained CA1 hippocampal neurons were obtained from coronal sections corresponding to Bregma −1.34 to −2.54 mm, and acquired using an Olympus IX83 P2ZF microscope with a 60X / 1.42 objective. The optical section thickness used was 0.56 µm. The criteria for selecting neurons and dendrites for analysis were as follows: stained neurons had to be isolated, fully impregnated, and present at least third-order branches for both apical and basal dendrites. Selected apical and basal dendrites must be of secondary order, non- bifurcating, and segments must be at least 10 µm long (15).

For spine density analysis, EFI projections of the stained neurons were obtained from Z-stack images using Olympus cellSens software. Spines were counted in at least 10 segments per animal. Thereafter, spine density was averaged per animal and results were expressed as the number of spines per µm of dendritic length (mean ± S.E.M.).

For spine morphology analysis, the Z-stack images were processed in Reconstruct software (ver. 1.1.0.0, http://synapses.clm.utexas.edu). The length and width of the spines within each 10 µm segment were traced. These two geometric parameters were used to assess morphological changes in dendrites. Because this study was conducted at the resolution limit of the optical microscope, spine length was measured from the base of the dendrite to the top of the spine head to minimize errors in spine neck tracing. Cumulative probability histograms of dendritic spine lengths and spine head widths were calculated.

For each experimental group, three mice per strain and sex were used. From each mouse, 10 dendritic segments (≥ 10 µm) from different neurons were analyzed for both apical and basal dendrites. For the morphological analysis, the total number of spines included was: WT male (n = 877), WT female (n = 717), KO male (n = 635), and KO female (n = 755) for apical dendrites, and WT male (n = 724), WT female (n = 661), KO male (n = 621), and KO female (n = 569) for basal dendrites.

**Transmission electron microscopy**

Lgals4-KO and control animals (8 months old) were anesthetized with sodium pentobarbital and transcardially perfused with saline followed by 4% PFA + 2%

glutaraldehyde in 0.1 M phosphate buffer (PB) at pH 7.4. Brains were post-fixed at RT for 2 h and overnight at 4°C, thoroughly washed in PB buffer, and coronal sections encompassing the dorsal CA1 region of the hippocampus (Bregma -1.34mm to -2.06mm) were obtained at 200 µm thickness using a vibrating microtome (Leica VT1200). Vibratome sections were postfixed with 1% osmium tetroxide and 0.8% aqueous potassium ferricyanide for 1h at 4 °C. After rinsing, samples were incubated with 0.15% tannic acid for 1 min at RT, 2% uranyl acetate for 1h at RT, dehydrated in a graded series of ethanol solutions at 4 °C, and embedded in epoxy resin (TAAB 812, TAAB Laboratories). Serial ultrathin sections (70 nm) were cut with an Ultracut E ultramicrotome (Leica), mounted on Formvar-carbon-coated Cu/Pd slot grids, and stained with 2% uranyl acetate and lead citrate. Samples were examined on a JEM-1400 Flash transmission electron microscope (Jeol, Japan) at 100 kV, with a Oneview 4K x 4K CMOS camera (Gatan, Pleasanton, CA USA).

Synaptic density was quantified in the *stratum radiatum* of hippocampal CA1 using the unbiased physical dissector method (35). Each dissector consisted of photomicrographs from the same position in two consecutive ultrathin sections, a reference and a look-up section. Synapses were identified by the presence of a postsynaptic density (PSD) and a presynaptic terminal containing synaptic vesicles, separated by a synaptic cleft. Perforated synapses were defined by the presence of a discontinuity in the PSD. Only synapses present in the reference section but absent in the look-up section, or vice versa, were manually counted by an observer blind to the genotype of the animal using ImageJ software. Synaptic density was calculated by dividing the number of synapses counted per dissector by the volume of the dissector (given by the frame area × dissector thickness). The dissector height was 140 nm. The total volume examined was 869.6 and 927.9 $\mu m^3$ for WT and Lgals4-KO groups, respectively; and the total number of synapses counted was 2143 and 2540 for WT and Lgals4-KO groups, respectively.

The lengths and areas of PSDs were quantified using a combination of manual annotation, deep learning, and image analysis software. A total of 90 micrographs were manually annotated using the "Training Labels" tool in OlyVia software (Olympus), where the PSD membrane, PSD area, and background regions were labelled. These annotated images served as the training set for a convolutional neural network (CNN) based on the U-Net architecture, implemented through Olympus' TruAI extension within the cellSens software (Olympus). The CNN training was configured with Balanced data sampling, Local Contrast normalization, Softmax cross-entropy loss, and the Adam optimizer. Data augmentation involved 90-degree rotations and mirroring to improve generalization. No additional image augmentation was applied. Training was conducted for 250,000 iterations with a similarity threshold of 0.71, using 10 of the 90 images for validation, a scaling factor of 25, and checkpoints saved every 6 epochs. Post-training, the model was applied to the remaining micrographs for automated detection and segmentation of PSDs. To ensure accuracy, every segmented image was reviewed by trained researchers blinded to genotype, and manually corrected in ImageJ when necessary to ensure precise PSD dimensions. Final measurements of PSD lengths and areas were obtained using a custom macro in ImageJ.

**Statistical analysis**

The following statistical tests were applied to the different data sets. Normal distribution of the data was determined using the Shapiro-Wilk test. For normally distributed data with equal variances, a two-tailed unpaired Student's t test was used to compare differences between two groups. For comparisons involving two factors, two-way ANOVA was applied. If normality could not be assumed, nonparametric tests were conducted: Mann Whitney U/Wilcoxon Rank Sum test for two groups and Kruskal-Wallis test for multiple group comparisons. Nonparametric multiple comparisons were done using Dunn´s test. For time-series data, two-way ANOVA with repeated measures was used. To compare distributions, the two-sample Kolmogorov-Smirnov test was applied. All data are presented as the mean ± S.E.M. Statistical significance is indicated as follows: $^{*}p < 0.05$; $^{**}p < 0.01$; $^{***}p < 0.001$.

**RNA sequencing and data analysis**

A total of 100 ng of total RNA was used to generate barcoded RNA-seq libraries using the NEBNext Ultra II Directional RNA Library Preparation Kit (New England Biolabs) according to manufacturer's instructions. First, poly(A)+ RNA was purified using poly-T oligo-attached magnetic beads, followed by fragmentation and synthesis of the first and second cDNA strands. Next, cDNA ends were repaired and adenylated, followed by NEBNext adaptor ligation, second strand removal, uracil excision from the adaptor, and

PCR amplification. Library size was assessed using the Agilent 2100 Bioanalyzer, and concentration was determined using the Qubit® fluorometer (Life Technologies). Libraries were sequenced at 2.5 nM on a HiSeq 4000 (Illumina) to generate 61 base single reads. FastQ files for each sample were obtained using bcl2fastq 2.20 Software (Illumina). RNA sequencing data analysis was conducted in R (version 4.4.0). Raw sequencing reads in FASTQ format were obtained from WT and KO samples, including both male and female groups. FastQC (Babraham Bioinformatics) was used for initial quality assessment, evaluating sequencing quality, adapter contamination, sequence duplication, and read length distribution. Adapter sequences were identified and removed using Trim Galore (Babraham Bioinformatics), followed by a second FastQC evaluation to ensure high-quality reads before alignment. The cleaned reads were aligned to the mm10 mouse genome (RefSeq GCF_000001635.27) using HISAT2 (version 2.1.1). Mapping quality was assessed using MultiQC (version 1.27), providing an overview of alignment efficiency across all samples. Gene-level quantification was performed with featureCounts function (version 2.0.8) (16), which assigned uniquely mapped reads to gene features using the NCBI RefSeq annotation. The resulting raw count matrix was used for subsequent differential expression analysis.

Differential expression analysis was conducted using DESeq2 (version 1.44.0) (17), which models RNA-seq count data with a negative binomial distribution to account for biological variability. Raw counts were imported into R and structured into a DESeqDataSet object which included genotype (WT vs. KO) and sex (male vs. female). Two statistical models were tested: a simple model (~ Strain + Sex), which assumes independent effects of genotype and sex, and a complex model (~ Strain + Sex + Strain:Sex), which includes an interaction term. A Likelihood Ratio Test (LRT) was performed to determine whether the interaction between strain and sex significantly influenced gene expression. Since no significant interaction effects were detected, the simpler model was selected for further analysis.

Normalization was performed using DESeq2's median of ratios method, which corrects for sequencing depth differences across samples. Hierarchical clustering analysis, implemented with pheatmap (version 1.0.12), confirmed biological consistency among replicates and grouped samples based on expression profiles.

To visualize differentially expressed genes (DEGs), a Volcano plot was generated using ggplot2 (version 3.5.1), displaying $\log_2$ fold change values against the adjusted p-values ($-\log_{10}P_{adj}$). Genes were classified as significantly differentially expressed if they met the criteria of $P_{adj} < 0.05$ and $|\log_2 \text{ fold change}| \geq 0.1$. To further explore alterations in synaptic gene expression, genes related to synaptic function were identified based on Gene Ontology (GO) annotations (GO.db version 3.19.1) and classified into four major categories. Presynaptic genes were identified using GO terms associated with neurotransmitter release, vesicle fusion, and calcium regulation, while postsynaptic genes were categorized based on GO annotations related to receptor clustering, synaptic scaffolding, and intracellular signaling at the postsynaptic membrane. Genes with shared synaptic functions, including those involved in synaptic signaling, organization, and trans-synaptic communication, were also identified, along with genes regulating synaptic plasticity, which play key roles in synaptic strength modulation, long-term potentiation, and depression. To visualize transcriptional differences in synaptic genes between WT and KO conditions, a heatmap was generated using ComplexHeatmap (version 2.20.0).

**Intestinal microbiota analysis**

Stool samples from 10 mice per strain (WT and Lgals4-KO), with equal numbers of males and females aged 6–7 months, were collected, and total DNA was extracted using the QIAamp Fast DNA Stool Mini Kit (Qiagen). The V3–V4 hypervariable region of the 16S rRNA gene was amplified via PCR, followed by the addition of sample-specific barcodes. Sequencing was performed on an Illumina MiSeq platform, generating 300 bp paired-end reads. Raw sequencing reads were processed using the DADA2 pipeline (v.1.32.0) in RStudio (v.2024.04.2+764, R v.4.4.0), including steps for quality filtering, trimming, dereplication, error correction, sequence variant inference, paired-end merging, and chimera removal. Taxonomic assignment was carried out using the SILVA database (version 138.1), with additional species-level identification through exact comparisons with SILVA and NCBI reference datasets. A phylogenetic tree was constructed, and the phyloseq package (v.1.48.0) in R was used for further analyses, including the removal of non-microbial taxa. Alpha diversity was assessed using observed richness, Shannon diversity, Faith's Phylogenetic Diversity, and Pielou's Evenness. Beta diversity was analyzed using Principal Coordinate Analysis (PCoA) based on Bray–Curtis dissimilarity,

Jaccard distance, and Unweighted UniFrac distance metrics. The significance of beta diversity differences between groups was tested using the adonis2 function (PERMANOVA) from the vegan package (v.2.6-6.1). Differential abundance of microbial taxa between experimental groups was evaluated using Microbiome differential abundance and correlation analyses with bias correction (ANCOM-BC, v.2.6.0) and Linear discriminant analysis Effect Size (LEfSe) from the microbiomeMarker package (v.1.10.0), which identified taxa with significant differences and estimated their effect sizes.

### Supplementary Figures

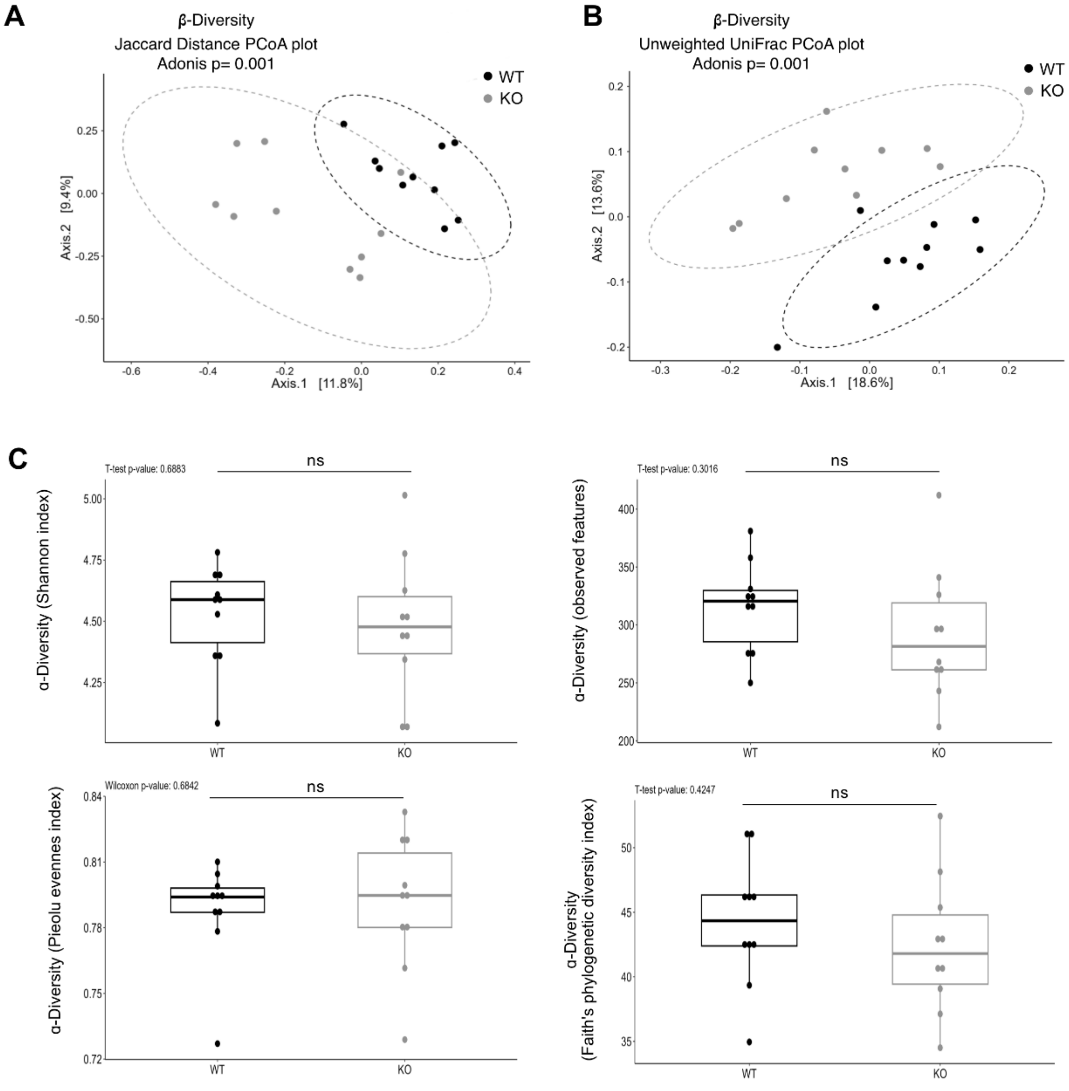


**Fig.S1**. Gut microbiota alpha and beta diversity in WT and Lgals4-KO mice. Principal coordinate analysis (PCoA) plots of bacterial beta diversity based on the Jaccard distance (A) and unweighted UniFrac distance (B) of gut microbiota samples from WT (black dots) and Lgals4-KO (grey dots) mice showing a significantly different sample distribution between groups (Adonis; p=0.001). Ellipses indicate 95% confidence interval, and the percentages in parentheses are the proportion of variation explained by the PCoA axis. (C) Box & Whisker representation of the intestinal microbiota alpha diversity for WT and Lgals4-KO mice, expressed by Shannon, observed features, Pielou evenness and faith's phylogenetic diversity indices. Unpaired t-test and Wilcoxon-rank sum test statistical analyses showed no significant differences between mice strains (ns). Animals used: WT n=10, KO n=10.

**A**

| Taxon | LFC | Adjusted p-value |
|---|---|---|
| g_Alistipes_s_timonensis | 3.5140 | 7.89E-06 |
| g_UCG-009 | -2.5371 | 0.00188 |
| g_Peptococcus | -2.8050 | 4.13E-10 |
| g_Paludicola | 1.7986 | 0.00306 |
| g_Prevotellaceae_s_NA | -3.6393 | 0.00745 |

**B**

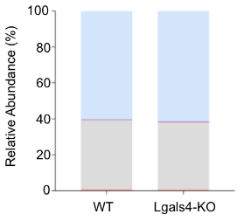


| Phylum | WT | Lgals4-KO |
|---|---|---|
| Actinobacteriota | 0.08 | 0.07 |
| Bacteroidota | 60.14 | 61.27 |
| Cyanobacteria | 0.00 | 0.01 |
| Desulfobacterota | 0.65 | 0.79 |
| Firmicutes | 38.34 | 37.19 |
| Proteobacteria | 0.79 | 0.67 |

**C**

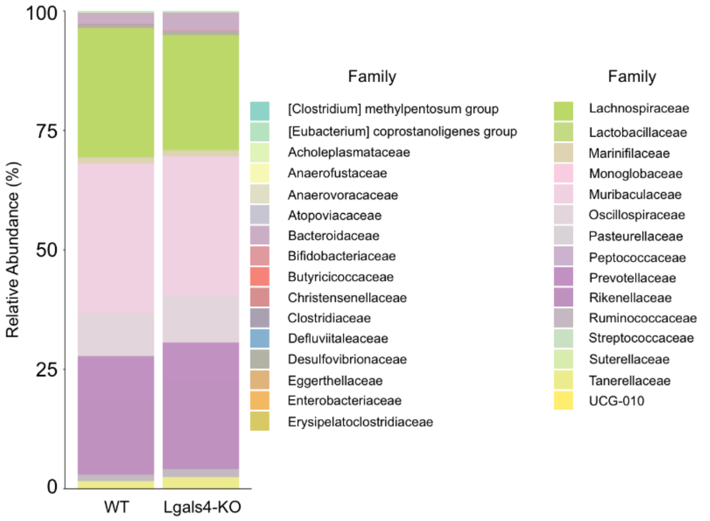


**Fig.S2.** Gut microbiota relative abundance in WT and Lgals4-KO mice. (A) Table shows the log fold change and adjusted p-values of differentially abundant taxa in the fecal microbiota of Lgals4-KO mice as determined by ANCOM-BC analysis. Stacked bar charts depicts the relative abundance of identified bacteria phyla (B) and families (C) in fecal microbiota samples from WT and Lgals4-KO mice.

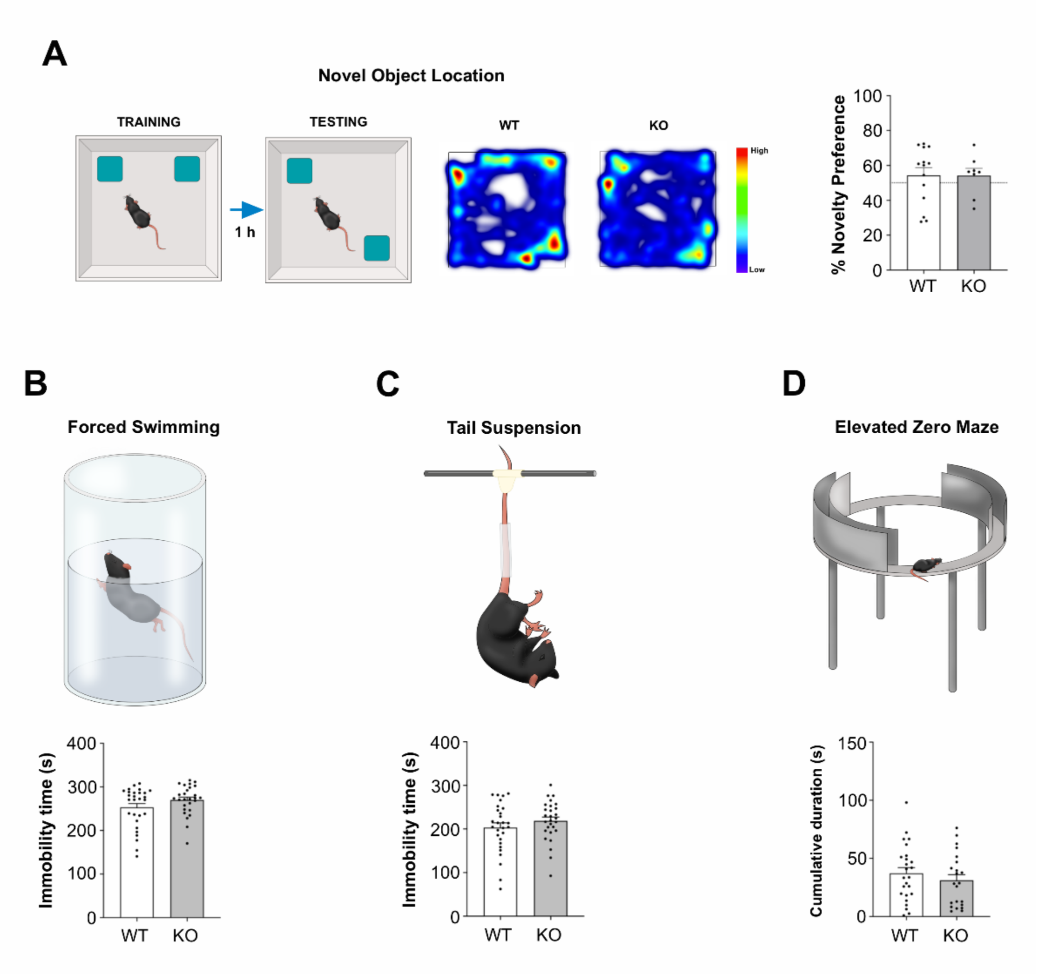

**Fig. S3**. Evaluation of memory and anxiety- and depression-like behaviors. (A) Schematic representation of the OL test and representative heatmaps illustrating the cumulative time mice spent exploring familiar and novel locations for WT (left) and KO (right) mice. Warm colors indicate more time spent exploring, while cold colors represent less. The bar graph shows the novelty preference percentage, with no significant difference between mouse strains (Mann-Whitney U test, p=0.8154; WT n=14, Lgals4-KO n=8). (B) Forced swim test. The top panel shows the test setup schematic, and the bar graph below displays total immobility time. No significant differences were observed between WT and Lgals4-KO mice (Mann-Whitney U test, p=0.4017; WT n=28, Lgals4-KO n=28). (C) Tail suspension test. Schematic representation of the test setup (top) and bar graph displaying the total immobility time (bottom). No significant differences were found between mice strains (Mann-Whitney U test, p=0.3162; WT n=28, Lgals4-KO n=28). (D) Elevated zero-maze test. Schematic representation (top) and bar graph displaying the time spent in open zones of the maze (bottom). No significant differences were observed (Unpaired t-test, p=0.3797; WT n=25, Lgals4-KO n=22). All graphs show the mean + SEM, with each dot representing an individual animal.

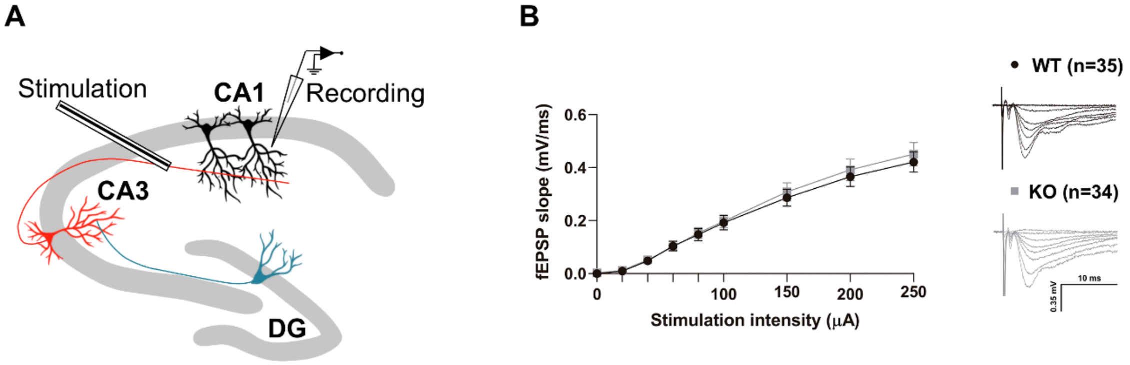


**Fig. S4.** Baseline input-output curves of evoked fEPSPs in the dorsal hippocampus. (A) Schematic diagram showing the in-vitro experimental setup of electrophysiological recordings from mice hippocampal brain slices. B) Input/output curves of averaged extracellular fEPSP slopes evoked at different stimulation intensities from WT and Lgals4-KO slices. Representative traces for each mice strain are shown on the right. The plot shows the mean slopes for each intensity used ± SEM. No significant differences were observed (Two-way repeated measures ANOVA; WT n=35 from 21 slices; KO n=34 from 20 slices; obtained from 10 animals per condition).

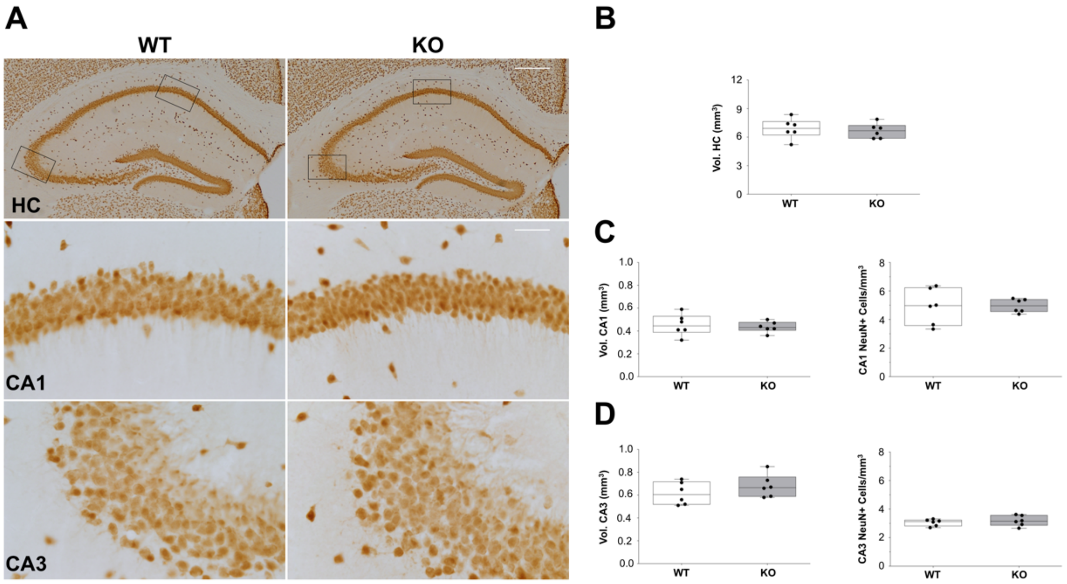


**Fig. S5.** Volume and neuronal density of the whole dorsal hippocampus, the CA1 and the CA3 regions in WT and Lgals4-KO mice. (A) Neuron-specific nuclear protein (NeuN) immunostaining of the hippocampus and magnifications within the CA1 and CA3 regions from top to bottom, respectively. HC scale bar = 400 µm. CA1 and CA3 scale bar = 40 µm. (B-D) Boxplots show the volume of the dorsal hippocampus (B), and the volume and neuronal density of the CA1 (C) and CA3 (D) regions. Cell density is expressed as the number of NeuN-positive cells (x$10^5$) per cubic millimeter. Dots on graphs represent the mean value for each animal. No significant differences were observed in any of the measured parameters (Unpaired t-test; WT n=6, Lgals4-KO n=6).

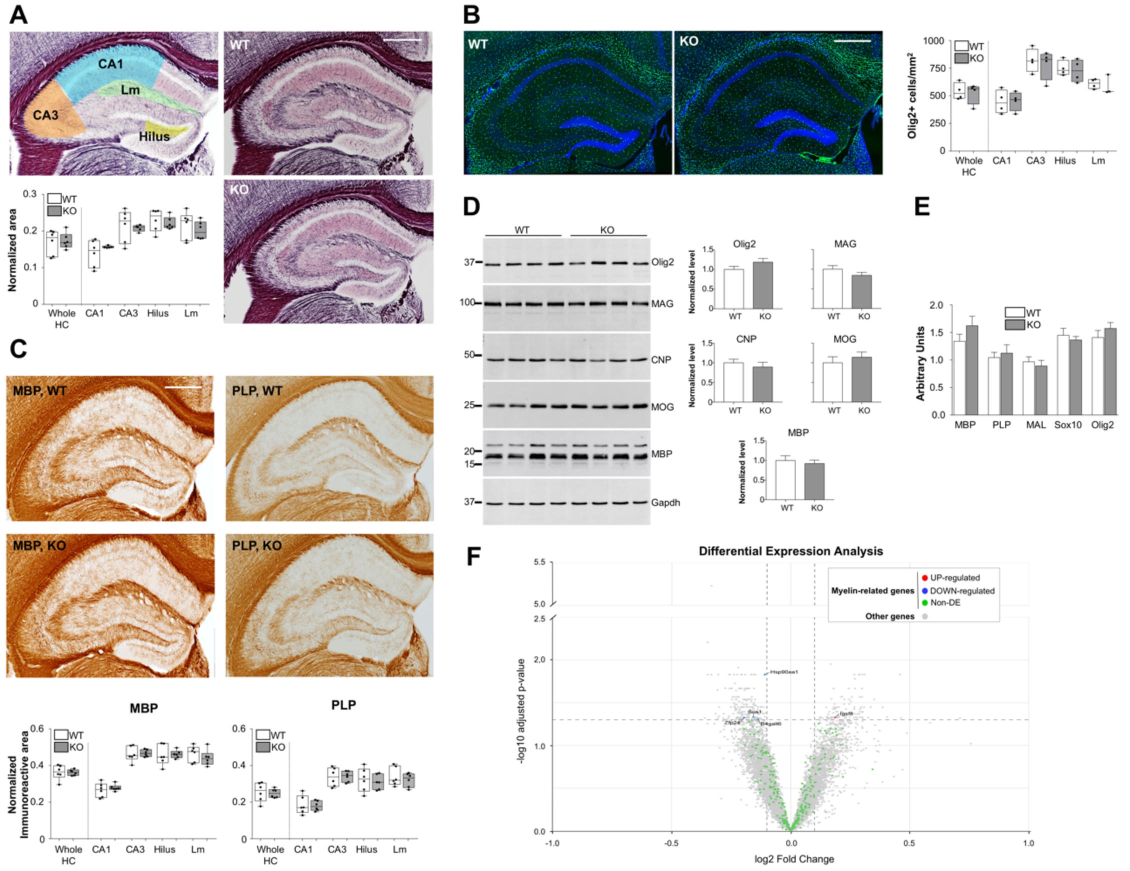


**Fig. S6** Unchanged distribution and expression of oligodendrocytes (OLG) and myelin markers in the hippocampus of Lgals4-KO mice. (A) The upper left panel shows the segmentation of CA1, CA3, *lacunosum moleculare* and hilus regions used for myelin analysis. The Black Gold staining (right panels) show the myelin distribution in WT (upper) and Lgals4-KO (lower) hippocampal slices. There were no significant differences in the normalized stained area between the two strains in the whole hippocampus (Whole HC) and its segmented areas (Mann Whitney U test; WT n=6, Lgals4-KO n=6). Scale bar: 500 µm. (B) Immunofluorescence staining for the oligodendrocyte marker Olig2 (green) in WT and Lgals4-KO hippocampal slices. The graph on the right shows no significant difference in the density of Olig2-expressing cells in the whole hippocampus, CA1, CA3, hilus and *lacunosum moleculare* (Lm) (Unpaired t-test; WT n=4, Lgals4-KO n=4). Scale bar: 500 µm. (C) Immunohistochemical staining for myelin markers MBP (left panels) and PLP1 (right panels) in hippocampal slices of WT and Lgals4-KO mice. The graphs on the bottom display the normalized area stained by each marker following the same segmentation as in (A). No significant differences were observed in any case (Unpaired t-test; WT n=6, Lgals4-KO n=6). Dots represent the mean value of all sections analyzed per animal. Scale bar: 500 µm. (D) Protein expression levels of selected myelin markers measured by WB. Hippocampal extracts from WT (left lanes) and Lgals4-KO (right lanes) animals were analyzed. GAPDH was used as loading control. In the graphs at the right, expressed values are means of normalized densitometric measures + SEM. No significant differences between WT and Lgals4-KO mice were observed (Unpaired t-test or Mann Whitney test when appropriate; WT n=4, Lgals4-KO n=4). (E) mRNA expression levels of mature-myelinating OLG markers *Mbp*, *Plp1* and *Mal*, and of OLG lineage markers *Olig2* and *Sox10* were measured by RT-PCR. No significant differences for any of the markers were observed when comparing WT to Lgals4-KO mice (Unpaired t-test; MBP: WT n=11, KO n=12; PLP1: WT n=11, KO n=11; MAL: WT n=10, KO n=11; Sox10: WT n=7, KO n=7; and Olig2: WT n=12, KO n=12). Values are means of normalized mRNA expression levels + SEM. (F) Volcano plot illustrates differentially expressed genes based on log2 fold change and adjusted p-values. Upregulated genes in Lgals4-KO mice relative to WT are shown in red, while downregulated genes are displayed in blue. Non-significant genes appear in gray. Genes related to myelin function, identified through Gene Ontology (GO) annotations, are highlighted in green.

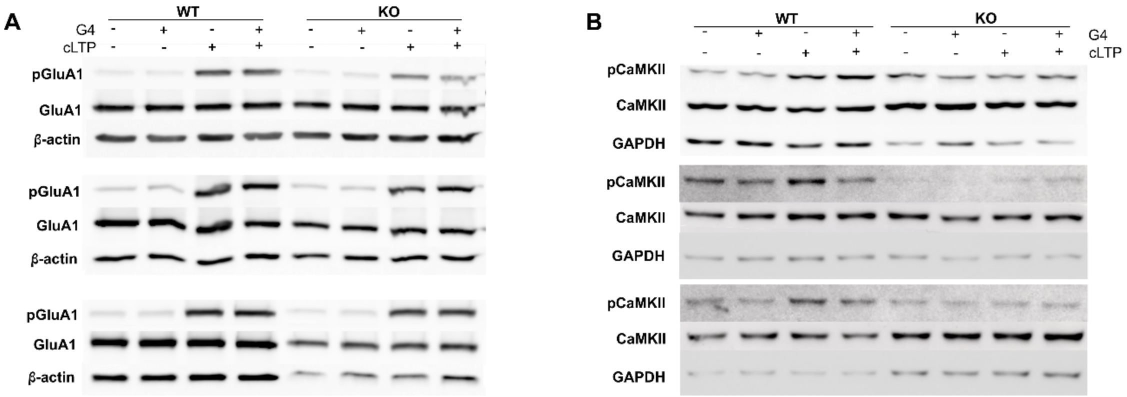


**Fig. S7**. Reduced activation of GluA1 receptors and αCaMKII after LTP induction in Lgals4-KO mice. Western blots of 3 independent experiments showing phosphorylated GluA1 (pGluA1; S845) (A), and phosphorylated αCaMKII (pCaMKII) (B) before (-) and after (+) cLTP induction in WT and Lgals4-KO hippocampal neuron lysates. Total GluA1 and ß-actin in (A), and total CaMKII and GAPDH in (B), are shown as loading controls and used for normalization.

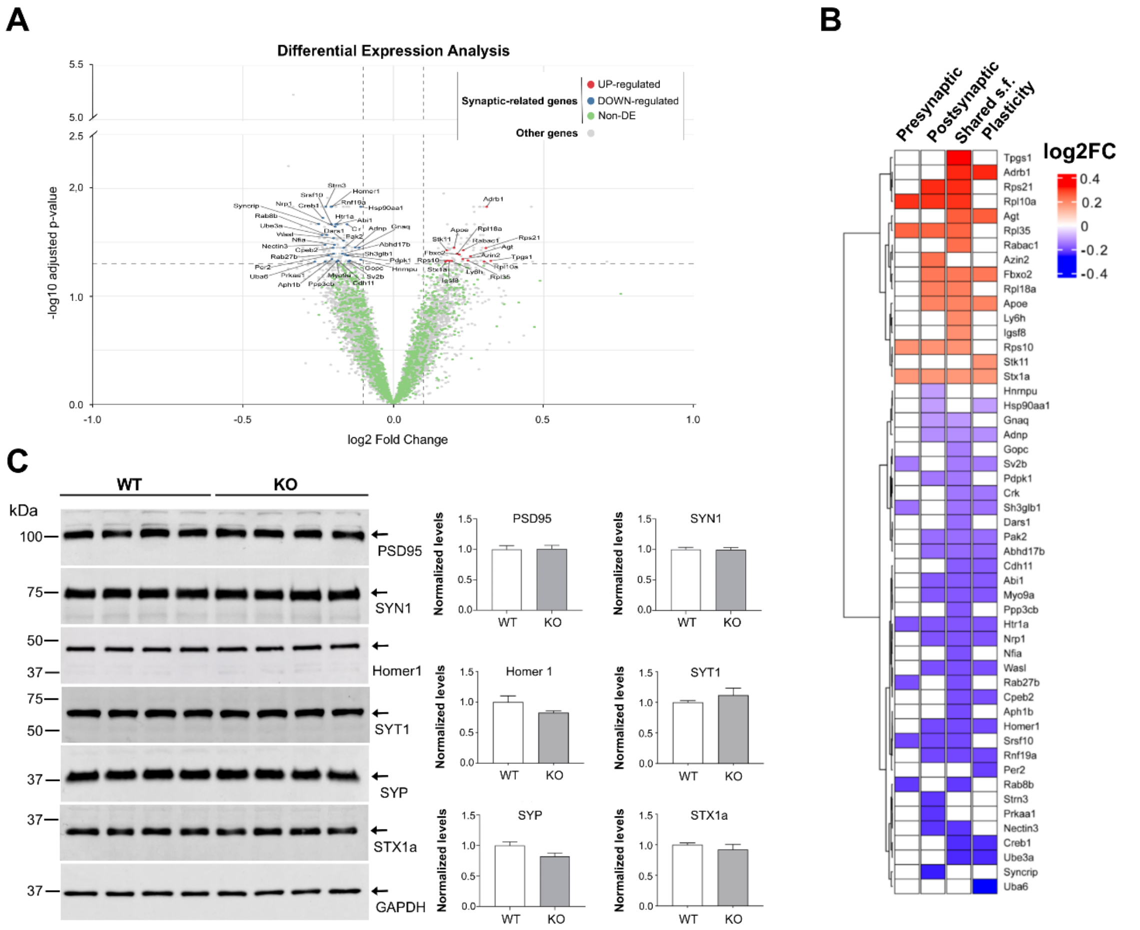


**Fig. S8** Differential expression of synaptic-related genes in Lgals4-KO mice. (A) Volcano plot illustrates differentially expressed genes based on log2 fold change and adjusted p-values. Upregulated genes in Lgals4-KO mice relative to WT are shown in red, while downregulated genes are displayed in blue. Non-significant genes appear in gray. Genes related to synaptic function, identified through Gene Ontology (GO) annotations, are highlighted in green. (B) Heatmap depicting the levels of differentially expressed genes involved in synaptic processes. The color scheme indicates the log2-transformed group mean of normalized counts. Rows categorize genes by synaptic location (Pre- or Postsynaptic), by their role in shared synaptic functions such as synaptic signaling, organization, and trans-synaptic communication (Shared s.f.)**,** or by their role in plasticity processes (Plasticity), based on selected Gene Ontology terms. (C) Protein expression levels of synaptic markers measured by WB. Left panels show representative WBs of hippocampal extracts from 4 WT and 4 Lgals4-KO animals, probed with antibodies against PSD-95, Synapsin 1 (SYN1), Homer 1, Synaptotagmin 1 (SYT1), Synaptophysin (SYP) and Syntaxin 1a (STX1a), and GAPDH as a loading control. The graphs on the right display the mean normalized densitometric measures + SEM. Quantitative analysis revealed trends in protein expression differences between WT and Lgals4-KO animals for Homer 1, SYT1 and SYP; however, none of these differences reached statistical significance (Unpaired t-test).

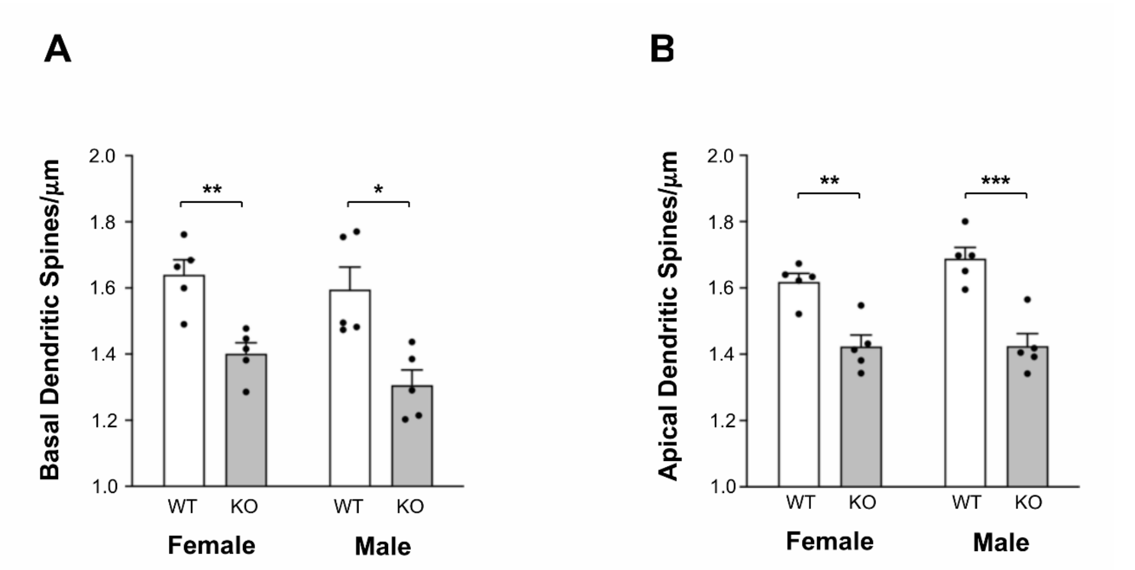


**Fig. S9** Dendritic spine density within the CA1 subfield according to animal sex. Both apical and basal dendritic spines exhibited significantly lower densities in female and male Lgals4-KO mice compared to their respective WT counterparts Basal dendrites: Two-way ANOVA revealed a significant effect of strain ($F(1,16) = 27.93$, $p<0.0001$), but not of sex ($F(1,16) = 1.975$, $p=0.1790$). Tukey's multiple comparisons test showed significant differences between WT and KO groups WT female vs. KO female, $p=0.0042$; WT male vs. KO male, $p=0.0180$. Apical dendrites: A significant effect of strain was also found ($F(1,16) = 47.85$, $p<0.0001$), with no significant effect of sex ($F(1,16) = 1.157$, $p=0.2980$). Tukey's test showed: WT female vs. KO female, $p=0.0037$; WT male vs. KO male, $p=0.0002$). The mean dendritic spine densities are shown. All data are expressed as means + SEM from 5 animals per strain and sex.

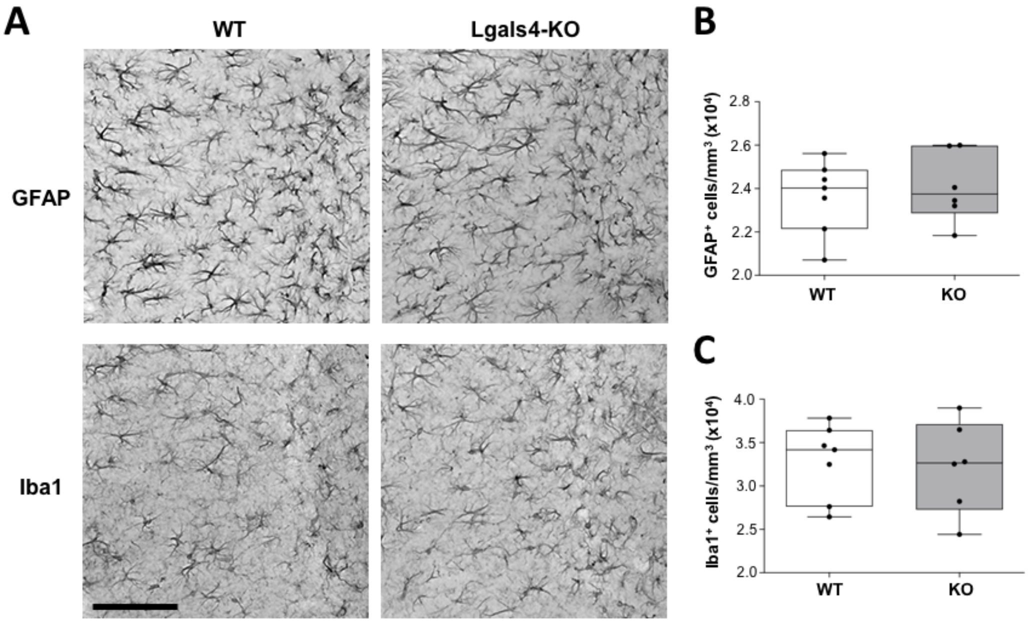


**Fig. S10**. Analysis of the density of astrocytes and microglia in the CA1 region of the hippocampus. (A) Representative images of GFAP (top) and Iba1 (bottom) staining in the *stratum radiatum* of the CA1 region of the hippocampus. Scale bar = 80 µm. (B-C) Quantification of the density of GFAP (B) and Iba1 (C) positive cells expressed as the number of cells ($x10^4$) per cubic millimeter. Data represented as a min-max boxplots show no significant differences between mice strains (Unpaired t-test, GFAP p=0.6259, Iba1 p=0.8381). Each dot represents the mean cell density per animal (WT n=7, Lgals4-KO n=6).

| | mRNA EXPRESSION (SNGLE CELL) | | | mRNA EXPRESSION (TISSUE) | | | | PROTEIN EXPRESSION (TISSUE) | | | |
|---|---|---|---|---|---|---|---|---|---|---|---|
| | | Lgals4 Expression Median of ln(1+CPM) | | | Lgals4 Expression Median of ln(1+TPM) | | Lgals4 Expression Median of ln(1+TPM) | | Galectin-4 Expression Parts per billion | | Galectin-4 Expression Parts per billion |
| BRAIN | Brain pericyte | 0 | 0 | Amygdala | 0,33 | brain | 1,61 | cerebellum | 35 | brain | 0,27 |
| | Endothelial cell | 0 | 0 | Anterior Cingulate Cortex | 0,30 | cerebellum | 1,95 | cerebral cortex | 23 | brain | 1,32 |
| | Astrocyte | 0 | 0 | Caudate | 0,39 | cerebral cortex | 1,95 | medulla oblongata | 13 | cerebral cortex | 15,8 |
| | Neuron | 0 | 0 | Cerebellar hemisphere | 1,04 | dentate gyrus | 1,95 | midbrain | 3 | midbrain | 17,8 |
| | Oligodendrocyte | 0 | 0 | Cerebellum | 1,11 | hippocampal formation | 1,10 | olfactory bulb | 24 | cerebellum | 17,6 |
| | Bergmann glial cell | 0 | 0 | Cortex | 0,38 | medial prefrontal cortex | 1,10 | | | cerebellum | 2,93 |
| | OPC | 0 | 0 | Frontal Cortex | 0,33 | olfactory bulb | 2,30 | | | medulla oblongata | 18,7 |
| | Microglial cell | 0 | 0 | Hippocampus | 0,35 | preoptic area | 1,39 | | | olfactory bulb | 43,1 |
| | | | | Hypothalamus | 0,41 | striatum | 1,61 | | | | |
| | | | | Nucleous Accumbens | 0,39 | subventricular zone | 2,08 | | | | |
| | | | | Putamen | 0,34 | | | | | | |
| | | | | Spinal Cord (Cervical C1) | 0,50 | | | | | | |
| | | | | Substantia nigra | 0,34 | | | | | | |
| INTESTINE | Enterocyte of epithelium | 8,49 | ND | Colon - Transverse | 7,19 | colon | 7,94 | colon | 7466 | colon | 3394 |
| | Globlet cell | 8,11 | ND | Terminal Ileum | 6,41 | | | ileum | 7936 | ileum | 3534 |
| | Epithelial cell | 7,66 | ND | | | | | | | jejunum | 3803 |
| | Brush cell of epithelium | 6,3 | ND | | | | | | | gut epithelium | 1105 |
| | Enteroendocrine cell | 5,73 | ND | | | | | | | | |
| | SOURCE | 1 | 2 | 3 | | 4 | | 5 | | 6 | |

| SOURCE | REPOSITORY | URL |
|---|---|---|
| 1 | Single Cell Expression Atlas | https://www.ebi.ac.uk/gxa/sc/home |
| 2 | Allen Brain Map | https://portal.brain-map.org/atlases-and-data/rnaseq/mouse-whole-cortex-and-hippocampus-10x |
| 3 | GTEx | https://gtexportal.org |
| 4 | Expression Atlas | https://www.ebi.ac.uk/gxa/genes/ |
| 5 | Expression Atlas | https://www.ebi.ac.uk/gxa/genes/ |
| 6 | PaxDb: Protein Abundance Database | https://pax-db.org/ |

**Fig. S11.** Overview of Gal-4 mRNA and protein expression in brain and intestine reported in different gene/protein expression repositories. mRNA expression in single cells and tissues, and protein expression in tissues are displayed. Note that Gal-4 expression is either extremely low or undetectable in brain tissues or cells, in agreement with the experimental results in Fig. S12.

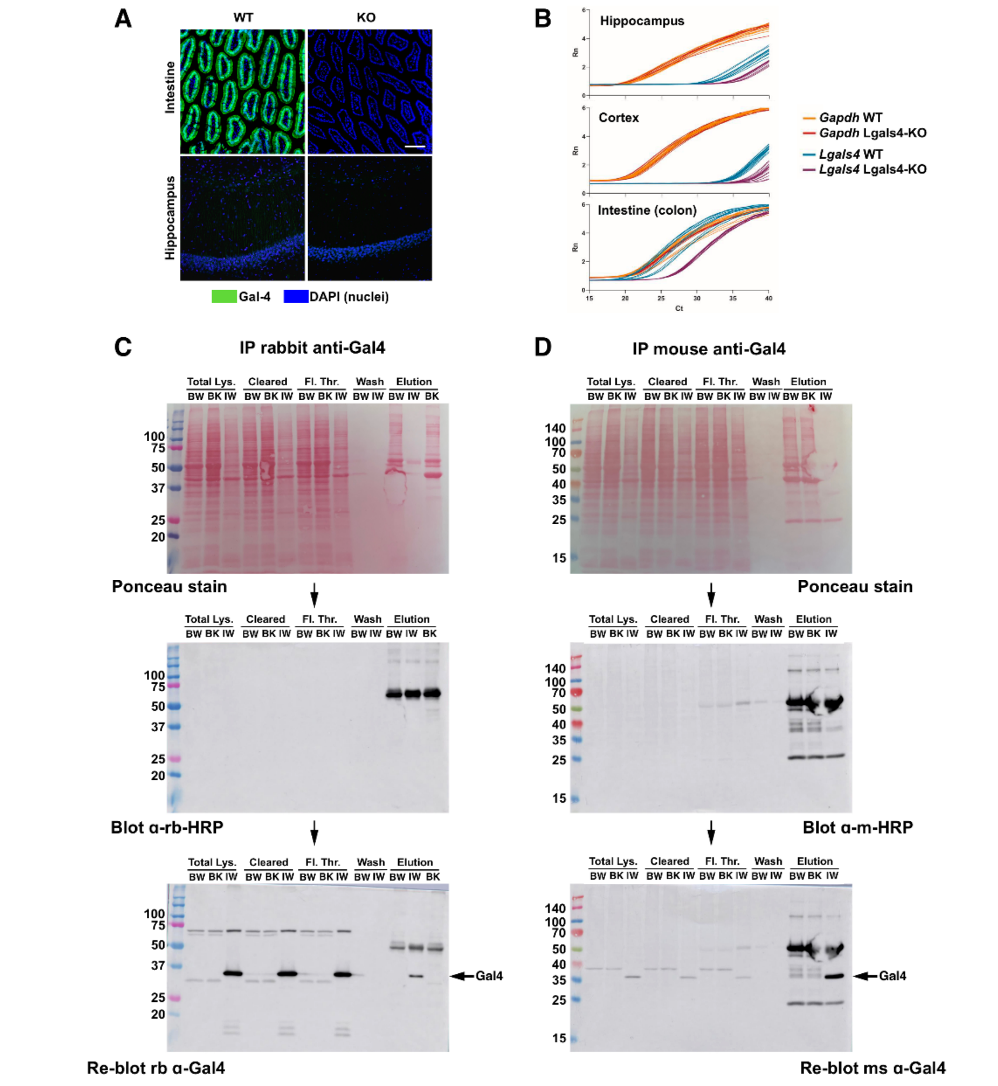


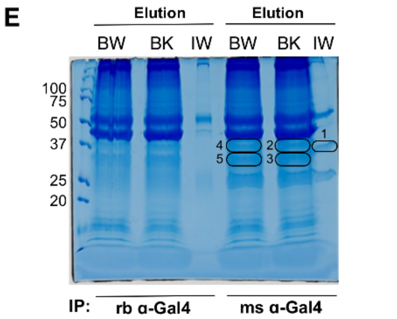


**F**

| Band | Accession | Score | Mass | matches | significant matches | sequences | significant sequences | Description |
|---|---|---|---|---|---|---|---|---|
| IW1 | A0A140LHI8 | 395 | 33979 | 21 | 14 | 15 | 11 | Galectin OS=Mus musculus GN=Lgals4 |
| IW1 | Q8C845 | 149 | 26784 | 7 | 7 | 7 | 7 | EF-hand domain-containing protein D2 |
| BK2 | A2AMW0 | 94 | 29562 | 2 | 2 | 2 | 2 | F-actin-capping protein subunit beta |
| BK2 | Q8C845 | 55 | 26784 | 2 | 2 | 2 | 2 | EF-hand domain-containing protein D2 |
| BK3 | Q8C845 | 255 | 26784 | 13 | 9 | 10 | 7 | EF-hand domain-containing protein D2 |
| BK3 | P47757 | 195 | 31611 | 13 | 10 | 10 | 8 | F-actin-capping protein subunit beta |
| BK3 | P14148 | 116 | 31457 | 3 | 2 | 3 | 2 | 60S ribosomal protein L7 |
| BK3 | P60710 | 115 | 42052 | 10 | 5 | 10 | 5 | Actin, cytoplasmic 1 |
| BK3 | Q8C2Q8 | 107 | 30294 | 6 | 6 | 6 | 6 | ATP synthase subunit |
| BW4 | P47754 | 129 | 33118 | 8 | 5 | 7 | 5 | F-actin-capping protein subunit alpha-2 |
| BW4 | P16125 | 116 | 36834 | 9 | 6 | 8 | 5 | L-lactate dehydrogenase B chain |
| BW5 | Q8C845 | 297 | 26784 | 13 | 10 | 12 | 9 | EF-hand domain-containing protein D2 |
| BW5 | P47757 | 179 | 31611 | 9 | 6 | 8 | 5 | F-actin-capping protein subunit beta |
| BW5 | P48962 | 109 | 33111 | 8 | 4 | 8 | 4 | ADP/ATP translocase 1 |
| BW5 | P60710 | 105 | 42052 | 8 | 4 | 7 | 3 | Actin, cytoplasmic 1 |
| BW5 | P62754 | 103 | 28834 | 5 | 3 | 4 | 3 | 40S ribosomal protein S6 |
| BW5 | P62908 | 101 | 26828 | 2 | 2 | 2 | 2 | 40S ribosomal protein S3 |

**Database: UniProt *MusMusculus***

**Fig. S12.** Gal-4 expression in intestinal and brain tissue. (A) Representative images of Gal-4 immunofluorescence labelling in intestinal and hippocampal tissue sections of WT and Lgals4-KO mice showing undetectable levels of Gal-4 protein in the hippocampus of WT mice. (Scale bar = 100 µm). (B) Gal-4 mRNA expression measured by RT-PCR in intestinal tissue of WT and LgalS4-KO mice. Graphs display the normalized fluorescence signal generated by the probes (Rn) vs the number of PCR cycles (Ct). The amplification of the endogenous control gene (*Gapdh*) is shown as orange and red lines, for the WT and Lgals4-KO, respectively. Blue and purple lines show the amplification of *Lgals4* gene in WT and Lgals4-KO samples, respectively. Note that in WT hippocampus and cortex (blue line) *Lgals4* is only detectable after 35 cycles, as it is for Lgals4-KO samples, indicating an extremely low expression of the *Lgals4* mRNA in these brain tissues. (C-D) Gal-4 immunoprecipitation from brain and intestinal tissues with rabbit-polyclonal (C) and mouse monoclonal (D) anti-galectin-4 antibodies. Ponceau staining (upper panels), no-primary antibody control blots with anti-species HRP-conjugated antibodies to detect non-specific bands (central panels), and re-blots with anti-Gal-4 antibodies (bottom panels) are shown. Gal-4 is detected exclusively by anti-Gal-4 WBs in immunoprecipitates from intestinal tissue. Tissue extracts were: brain wild type (BW), brain Lgals4-KO (BK), and intestine wild type (IW). Loaded samples were: total lysate (Total Lys.), cleared total lysate (cleared), protein A/G column flow-through (Fl. Thr.), column wash (Wash) and column elution (Elution). (E) Coomassie-stained PAGE-SDS gel of brain and intestine extracts immunoprecipitated with anti-Gal-4 antibodies. Gel bands split off for proteomic analysis are numbered 1-5. (F) Identification of the proteins in gel bands obtained in (E) after in-gel digestion and LC-MS/MS analysis. The table displays the best matched annotations (highest scores) in each gel band. Note that Gal-4 (Lgals4) is identified in WT intestine (IW1, highlighted in yellow), while it is not detected in WT brain (BW4 and BW5).

### Supplementary Tables

| Antigen | Host | Clone | Cat. No. | Source | Working dilution | |
|---|---|---|---|---|---|---|
| | | | | | IHQ/IF/IC | WB |
| **CaMKII** | Mouse | 6G9 | MA 1-048 | ThermoFisher | | 1/1000 |
| **CaMKII-P-T286** | Rabbit | Polyclonal | Ab5683 | Abcam | 1/300 | 1/1000 |
| **CNPasa** | Rabbit | D83E10 | 5664 | Cell Signaling | | 1/1000 |
| **Galectin-4** | Goat | Polyclonal | AF1227 | RD Systems | 1/100 | 1/1000 |
| **Galectin-4** | Rabbit | Polyclonal | -- | Gabius H.J. lab | 1/100 | 1/1000 |
| **Galectin-4** | Mouse | B-9 | Sc-271533 | Santa Cruz | 1/100 | 1/1000 |
| **GAPDH** | Mouse | 6C5 | MAB374 | Merck Millipore | | 1/10000 |
| **GFAP** | Mouse | GA5 | 3670 | Cell Signaling | 1/400 | |
| **GluA1-CT** | Rabbit | Polyclonal | Ab31232 | Abcam | 1/100 | |
| **GluA1-NT** | Mouse | RH95 | MAB2263 | Merck Millipore | 1/100 | 1/1000 |
| **GluA1-P-S845** | Rabbit | EPR2148 | Ab76321 | Abcam | | 1/1000 |
| **Homer 1** | Rabbit | Polyclonal | PA 5-21487 | ThermoFisher | | 1/2000 |
| **Iba1** | Rabbit | Polyclonal | 10904-1-AP | Proteintech | 1/400 | |
| **MAG** | Goat | Polyclonal | Sc-9544 | Santa Cruz | | 1/1000 |
| **MBP** | Rat | 12 | Ab7349-1 | Abcam | 1/200 | 1/1000 |
| **MOG** | Rabbit | E5K6T | 96457 | Cell Signaling | | 1/1000 |
| **NeuN** | Rabbit | D3S3I | 12943 | Cell Signaling | 1/700 | |
| **nNOS** | Goat | Polyclonal | ab1376 | Abcam | | 1/1000 |
| **Olig2** | Mouse | 211F1.1 | MABN50 | Merck Millipore | 1/500 | 1/2000 |
| **PLP1** | Rabbit | E9V1 | 28702S | Cell Signaling | 1/200 | |
| **PSD95** | Rabbit | Polyclonal | Ab18258 | Abcam | | 1/1000 |
| **STX1a** | Mouse | C-5 | Sc-17836 | Santa Cruz | | 1/1000 |
| **SYN1** | Rabbit | Polyclonal | ANR-014 | Alomone | | 1/1000 |
| **SYP** | Rabbit | Polyclonal | ANR-013 | Alomone | | 1/1000 |
| **SYT1** | Rabbit | Polyclonal | ANR-003 | Alomone | | 1/1000 |
| **Rabbit IgG - Alexa Fluor 488** | Donkey | | A21206 | Invitrogen | 1/500 | |
| **Rabbit IgG-Alexa Fluor 594** | Donkey | | A21207 | Invitrogen | 1/500 | |
| **Mouse IgG - Alexa Fluor 488** | Donkey | | A21202 | Invitrogen | 1/500 | |
| **Rabbit IgG-biotin** | Goat | | BA-1000 | Vector | 1/500 | |
| **Rat IgG-biotin** | Goat | | 629540 | Zymed | 1/500 | |
| **Mouse IgG-biotin** | Horse | | BA-2000 | Vector | 1/500 | |
| **Goat IgG HRP** | Rabbit | | A8919 | Sigma | | 1/5000 |
| **Rabbit IgG HRP** | Donkey | | NA-934V | GE Healthcare | | 1/5000 |
| **Rat IgG HRP** | Rabbit | | A9542 | Sigma | | 1/5000 |
| **Mouse IgG HRP** | Sheep | | NA931V | GE Healthcare | | 1/5000 |

**Table S1**. Antibodies used in this work: Sources and working dilutions.

| Gene | Reference |
|---|---|
| Lgals4 G07 | Mm01179060_m1 |
| SOX10 | Mm00569909_m1 |
| MAL | Mm01339780_m1 |
| MBP | Mm01266402_m1 |
| Olig2 | Mm01210556_m1 |
| PLP1 | Mm01297210_m1 |
| GAPDH | Mm99999915_g1 |
| β-Actin | Mm02619580_g1 |

**Table S2**. TaqMan probes (Applied Biosystems) used in expression measurements by RT-PCR. All probes were conjugated to fluorescein amidite (FAM) fluorophore.
